# Organ-specific NLR resistance gene expression varies with plant symbiotic status

**DOI:** 10.1101/135764

**Authors:** David Munch, Vikas Gupta, Asger Bachmann, Wolfgang Busch, Simon Kelly, Terry Mun, Stig Uggerhøj Andersen

**Author notes:** Author for correspondence, Stig Uggerhøj Andersen,. **Abbreviations:** *Arabidopsis*, AM, CC, CNL, ETI, *Lotus*, LRR, MAMP, *Medicago*, NB-ARC, NBD, NF, NLR, PRRs, RNL, TIR, TNL, XNL.

## Abstract

Nucleotide-binding site leucine-rich repeat resistance genes (NLRs) allow plants to detect microbial effectors. We hypothesized that NLR expression patterns would reflect organ-specific differences in effector challenge and tested this by carrying out a meta-analysis of expression data for 1,235 NLRs from 9 plant species. We found stable NLR root/shoot expression ratios within species, suggesting organ-specific hardwiring of NLR expression patterns in anticipation of distinct challenges. Most monocot and dicot plant species preferentially expressed NLRs in roots. In contrast, *Brassicaceae* species, including oilseed rape and the model plant *Arabidopsis thaliana*, were unique in showing NLR expression skewed towards the shoot across multiple phylogenetically distinct groups of NLRs. The *Brassicaceae* NLR expression shift coincides with loss of the endomycorrhization pathway, which enables intracellular root infection by symbionts. We propose that its loss offer two likely explanations for the unusual *Brassicaceae* NLR expression pattern: loss of NLR-guarded symbiotic components and elimination of constraints on general root defences associated with exempting symbionts from targeting. This hypothesis is consistent with the existence of *Brassicaceae*-specific receptors for conserved microbial molecules and suggests that *Brassicaceae* species are rich sources of unique antimicrobial root defences.

## Introduction

The sessile nature of vascular plants has spurred development of mechanisms for coping with biotic and abiotic stresses and for optimizing uptake of inorganic compounds under low nutrient availability. In response to these challenges, plant roots and shoots have evolved specialized functions above and below ground, where they have also adapted to interact with the distinct microbial communities of the phyllo- or rhizosphere. These diverse plant-microbe interactions range from symbiosis over parasitism to pathogenic infection (Bulgarelli et al. 2013; Fatima et al. 2015; Vandenkoornhuyse et al. 2015).

Reflecting the different characteristics of plant roots and shoots, distinct host-microbe combinations have been used to unravel the molecular components required for trans-species interaction and communication. In plant shoots, the focus has almost exclusively been on pathogenic interactions, where work in the model plant *Arabidopsis thaliana* (*Arabidopsis*) from the Brassicaceae family has provided great insight into plant immunity (Jones and Dangl 2006; Nishimura et al. 2010). Passive defences, such as the waxy cuticle on epidermal cells, cell walls and preformed anti-microbial chemicals form the first barriers for microbes and are often sufficient for deterring would-be pathogens (Thordal-Christensen 2003). Microbes that successfully evade these obstacles encounter a large repertoire of resistance (R) proteins in the form of trans-membrane receptor-like proteins and receptor-like kinases on the surface of plant cells, which recognize conserved microbe-associated molecular patterns (MAMPs). Upon activation, these pattern-recognition receptors (PRRs) trigger complex intracellular signalling cascades, such as phytohormone perturbations, accumulation of ions, mitogen-activated protein kinase activation and production of reactive oxygen species, ultimately leading to transcriptional and translational changes that promote the production of defence compounds ( Pel et al. 2012; Muthamilarasan et al. 2013).

To escape this MAMP-triggered immunity (MTI), microbes have evolved effectors that are injected into the plant cell cytoplasm using specialized secretion systems that penetrate the plant cell membrane. Upon translocation, these effectors target components of the defence machinery, suppressing immune signalling and gene expression through degradation, allosteric or covalent modification of host molecules, thus adapting the local environment to be more suitable for microbial growth and improving the chances of successful tissue colonization (Jones and Dangl 2006; Xin et al. 2013; Le Fevre et al. 2015). In response, plant cells employ a family of intracellular R proteins that recognize effectors either by direct interaction, or indirectly through detection of modifications made to host proteins (Khan et al. 2015). Effector-triggered immunity (ETI) activation by an intracellular R protein leads to a stronger immune response than that of MTI and is often associated with localized cell death to limit the spread of biotrophic pathogens (Jones and Dangl 2006; Hofius et al. 2007).

The majority of intracellular R proteins share a similar structure with an amino-terminal signalling domain, followed by a highly conserved nucleotide binding domain (NBD) and a carboxy-terminal leucine-rich repeat (LRR) domain of variable length ( van der Biezen et al. 1998; Takken et al. 2012). This class of R proteins are referred to as nucleotide-binding site leucine-rich repeat (NLR) proteins. The NBD domain class is shared by Apaf1, plant R proteins and CED4 (NB-ARC) and is highly conserved among all NLR proteins. It acts as a molecular switch, and cycles between active ATP-bound and inactive ADP-bound states depending on the activity of the LRR domain. The LRR domain is believed to be directly involved in protein-protein interactions with microbial effectors or host proteins and to function by auto-suppressing the NBD domain of the NLR (Jones and Jones 1997; Takken et al. 2006; Marquenet et al. 2007; Lukasik et al. 2009; Takken et al. 2012). The amino-terminal signalling domain is generally divided into two separate classes based on homology to either the signalling domain of Toll/Interleukin-1 Receptors (TIR) or the presence of a coiled-coil (CC) domain. These two distinct signalling components share common downstream signalling pathways, however both classes have also been observed to activate separate downstream components (Aarts et al. 1998; Falk et al. 1999; Meyers et al. 1999; Pan et al. 2000; Takken et al. 2006; Hofius et al. 2009). While both CC and TIR type NLRs (CNLs and TNLs, respectively) are widely distributed in dicots, canonical TNLs appear to be absent in monocots ( Meyers et al. 1999; Pan et al. 2000; Meyers et al. 2002; Tarr et al. 2009). In addition, variations of the signalling domain-NBD-LRR (NLR) structure can be found in most plant species, with NBD-containing proteins lacking either the amino-terminal signalling domain or the carboxy-terminal LRR domain, or having juxtaposed non-canonical domains, extending their flexibility as signalling components or effector decoys for host proteins (Bonardi et al. 2012; Kroj et al. 2016).

Whilst many NLRs play important roles in *Arabidopsis* shoot immunity, little is known about how *Arabidopsis* roots mount immune response against microbes, or what role NLRs play. However, the PRR FLAGELLIN-SENSITIVE2 is fully functional in roots and activates similar downstream MAP-kinase cascades in both root and shoot (Millet et al. 2010). There are reported differences between roots and shoots for the phytohormone salicylic acid, which is considered a requirement for basal defence in leaves against biotrophic pathogens, but does not appear to be as important in root immune responses (Jones and Dangl 2006; Millet et al. 2010).

Unlike the work on *Arabidopsis* pathogen responses, studies of root-microbe interactions have focused on endosymbiosis. Up to 90% of all terrestrial plants are believed to associate with arbuscular mycorrhizal (AM) fungi to enhance their acquisition of phosphorus and other nutrients. Plant associations with nitrogen-fixing bacteria contained within nodules is restricted to around 10 families, including the agriculturally important Fabaceae (legume) family (Doyle 1998; Gualtieri et al. 2000; Parniske 2008). *Arabidopsis* belongs to the *Brassicaceae* family which is one of the few plant families that has lost the capacity for root endosymbiosis with mycorrhizal fungi that is ancestral to the Angiospermae (flowering plants) (Gualtieri et al. 2000; Smith et al. 2010; Delaux et al. 2014). Two model plants from the legume family, *Lotus japonicus* (*Lotus*) and *Medicago truncatula* (*Medicago*), have been extensively studied for unravelling the genetic pathways required for root nodulation through their symbiotic association with gram-negative soil bacteria collectively referred to as rhizobia (Barker et al. 1990; Handberg and Stougaard 1992). This work has led to the discovery of nodulation factors (NF), a key signal molecule secreted by rhizobia, and several host receptors that perceive and transduce the signal through regulatory components to modulate downstream transcriptional regulation and coordinate nodule organogenesis and infection of these by nitrogen-fixing rhizobia (Long 1989; Schauser et al. 1999; Limpens et al. 2003; E. B. Madsen et al. 2003; Radutoiu et al. 2003; Lévy et al. 2004; Kalo et al. 2005; Smit et al. 2005; Tirichine et al. 2006; Kouchi et al. 2010; Madsen et al. 2010). Similar to NF produced by rhizobia, AM fungi secrete Myc factors to activate symbiotic signalling in the host. Despite their distinct phenotypic characteristics, AM and nodulation pathways share conserved genetic components, likely owing to their common evolutionary origin (Oldroyd and Downie 2006; Parniske 2008; Banba et al. 2008; Singh and Parniske 2012; Guillotin, Couzigou, and Combier 2016).

Despite the history of focusing on pathogenic plant-microbe interactions in plant shoots and on symbiotic interactions in roots, both organs are prone to pathogen infection and would presumably be protected by NLR proteins present in cells subject to effector challenge. Currently, little is known about the expression characteristics of NLRs and, unless they are ubiquitously expressed across all plant organs, NLR gene expression patterns could provide indications about differences in pathogen effector pressures between plant tissues and across plant species. Here we present a meta-analysis of NLR gene expression data, including plant species with and without the capacity for mycorrhizal and/or root nodule symbiosis. The analysis revealed stable root to shoot NLR gene expression ratios within species, with all of the endomycorrhizal plant species examined predominantly expressing NLRs in roots. In contrast, large differences were found between species, with the Brassicaceae family displaying an aberrant shoot-skewed expression, which suggested an unusual mode of plant-microbe interaction for this plant family.

## Results

### NLR gene expression varies between tissues in a species-specific manner

Individual plant organs have evolved to function in specific environments, where they interact with distinct microbiota (Vandenkoornhuyse et al. 2015). To investigate if NLR expression patterns reflected these tissue differences we identified all putative NLRs in *Lotus* and *Arabidopsis*, where expression atlas data was available for multiple tissues (Schmid et al. 2005; Høgslund et al. 2009; Verdier et al. 2013) (Supplemental table 1-2 **and Supplemental file 1**). We then examined the available expression data and identified genes predominantly expressed in reproductive, shoot, root or root nodule tissues. NLR expression in *Lotus* shoot and nodule tissues did not show significant differences compared to overall gene expression, but reproductive tissues showed strong depletion of NLR expression and *Lotus* roots displayed a significant enrichment of NLR expression (Figure 1A-B). For *Arabidopsis*, reproductive tissues also showed a significant depletion of expressed NLR genes, but *Arabidopsis* roots did not show enriched NLR gene expression. Instead, *Arabidopsis* shoots displayed a significant enrichment of NLR gene expression (Figure 1C-D).

**Figure 1.**
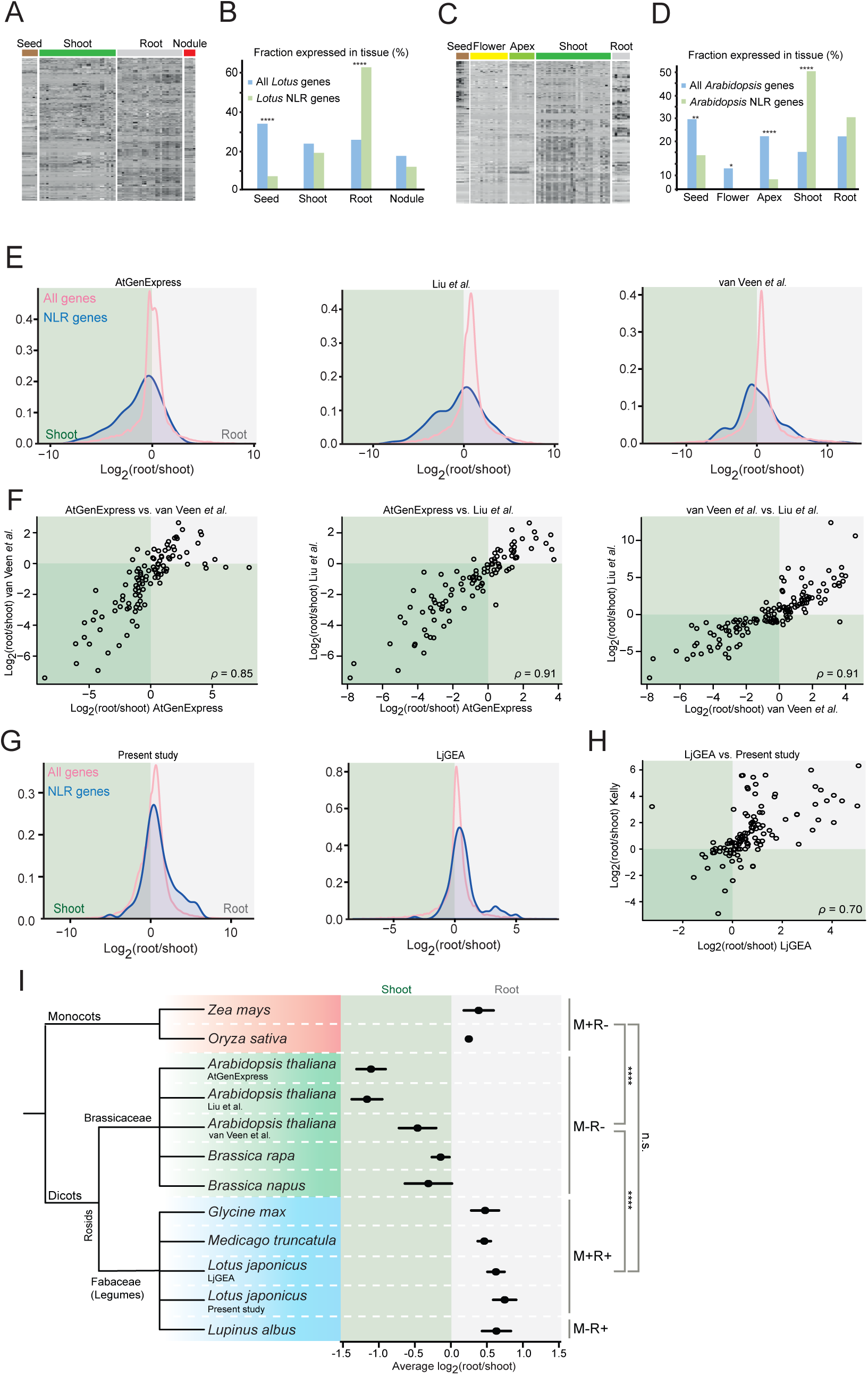
NLR gene expression patterns. **A)** Expression patterns of 198 putative *Lotus* NLR genes. **B)** Enrichment of *Lotus* NLR genes by tissue type. The fraction of all genes and NLR genes with enriched expression in the given tissue are shown. **C)** Expression patterns of 160 putative *Arabidopsis* NLR genes. **D)** Enrichment of *Arabidopsis* NLR genes by tissue type. The fraction of all genes and NLR genes with enriched expression in the given tissue are shown. *P*-values indicate the probability that the fraction of NLR genes showing enriched expression in a specific tissue is identical to that of all genes. **E)** Density plots displaying the distribution of the logarithm of the root/shoot expression ratios of all genes and NLR genes for each *Arabidopsis* expression data set indicated. See main text for sources. **F)** Root/shoot expression correlations for *Arabidopsis* NLR genes from the three data sets shown in Figure 1E. Each circle represents one NLR gene for which expression data is available in both of the datasets compared. **G)** Density plots displaying the distribution of the logarithm of the root/shoot expression ratios of all genes and NLR genes, for each *Lotus* expression data set indicated. **H)** Root/shoot expression correlations for *Lotus* NLR genes between two datasets. Each circle represents one NLR gene for which expression data is available in both datasets. **I)** Phylogenetic tree of species for which both shoot and root expression data is available, along with their average NLR gene root/shoot expression values (black dots). Error bars indicate SEM. The symbiotic status of each species is indicated on the right; M: Mycorrhiza. R: Rhizobia. +: engages in endosymbiosis. -: does not engage in endosymbiosis. Significance of each species group is indicated on the far right; ****: Significant difference with p ≤ 0.0001. n.s.: No significance. ANOVA and Tukey’s multiple comparison test was used for calculation of *P*-values. See Supplemental table 5 for *P*-values for inter-group and inter-species differences.

To investigate if the contrasting root/shoot NLR gene expression ratios were general for the two species, we examined additional data sets. For *Arabidopsis*, we quantified NLR root/shoot expression ratios based on two recent RNA-seq experiments including both root and shoot samples in the same experimental series (van Veen et al. 2016; Liu et al. 2016). Both RNA-seq data sets showed a clear shoot skew for *Arabidopsis* NLRs relative to the average expression ratio for all genes (Figure 1E), and the NLR expression ratios were strongly correlated across array and RNA-seq experiments (Figure 1F). Since no equivalent data sets were available for *Lotus*, we carried out an RNA-seq experiment including mock and rhizobium inoculated root and shoot samples. For *Lotus*, the RNA-seq data was also consistent with the array data in showing a pronounced root skewed NLR expression (Figure 1G-H). Since bacterial inoculation could potentially influence NLR root/shoot expression ratios, we compared *Lotus* inoculated and uninoculated samples, but found no significant differences in the root/shoot NLR expression ratios for neither the array nor the RNA-seq experiment (Supplemental figure 1). NLR root/shoot expression ratios thus showed clear differences between *Lotus* and *Arabidopsis*, and these differences were consistent across independent experiments carried out using either array or RNA-seq methodology for transcript quantification, indicating that regulation of NLR gene expression varied between organs in a species-specific manner.

### The *Brassicaceae* family shows aberrant shoot-skewed NLR gene expression

To determine which of these contrasting patterns of NLR gene expression was predominant among flowering plants, we analysed additional species for which root and shoot tissues had been subjected to global expression profiling in the same experiment. These included three legume species (*Medicago*, *Glycine max*, *Lupinus albus*), two *Brassicaceae* family members (*Brassica rapa* ssp. *pekinensis*, *Brassica napus*) and two monocots (*Zea mays, Oryza sativa*) (Figure 1I and Supplemental figure 2). We calculated root/shoot expression ratios for whole transcriptomes, including only samples where root and shoot tissues had been analysed in the same experimental series (Supplemental tables 1-2). We identified a total of 2,167 NLR genes across the selected species, and expression data was available for 1,235 out of the 2,167 NLRs (Supplemental table 3). Like *Lotus*, the three other dicot legumes and the two monocots displayed NLR gene expression skewed towards the root when compared to the overall gene expression pattern (Figure 1I, Supplemental figure 2 and Supplemental table 4). In comparison, the three *Brassicaceae* species stood out by displaying shoot-skewed NLR gene expression (Figure 1I and Supplemental table 4). Comparisons within either the legume, *Brassicaceae* or monocot groups did not show any statistically significant differences. However, when we compared between species groups, many comparisons showed significant differences, with the differences between *Brassicaceae* versus both legumes and monocots highly significant (Figure 1I and Supplemental table 5). Among the flowering plants investigated, shoot-skewed expression of NLR genes was a feature exclusive to the dicot *Brassicaceae* family, while the remaining monocots and dicot species all displayed root-skewed expression.

### The *Brassicaceae* expression shift is seen across multiple NLR clades

We speculated if the *Brassicaceae* expression shift could have been caused by the loss of a specialized set of phylogenetically related NLRs evolved specifically to guard the root endosymbiotic machinery or other root specific pathways. To test this hypothesis, we categorized all identified NLRs by aligning their NBDs and constructing a phylogenetic tree based on 2,033 sequences (Figure 2A). In addition to the previously mentioned species, we included the carnivorous and submerged aquatic bladderwort *Utricularia gibba* from the Asterids clade, which lacks a true root (Ibarra-Laclette et al. 2013). The phylogenetic analysis allowed us to identify five well-supported major NLR clades (Figure 2B, Supplemental file 2, and Supplemental table 7). We also categorized the NLRs based on the presence of TIR, CC or CC_R_ amino terminal signalling domains (Xiao et al. 2001; Meyers et al. 2003; Shao et al. 2016) and compared these results to our phylogenetic analysis (Supplemental figure 3 and Supplemental table 6). Hereafter, we refer to NLRs containing TIR, CC and CC_R_ domains as TNLs, CNLs and RNLs, respectively. NLRs containing neither of the three described domains are referred to as XNLs. Clade 1 was highly enriched in TNLs (708/806), CNLs dominated clade 2 (307/510) and clade 4 (326/385), clade 5 was enriched for RNLs (62/87), and clade 3 contained mainly XNLs (211/245) (Supplemental table 7). The clear correlation between domain structure and the NBD-based phylogeny indicated that the NBD sequences contained sufficient information for inferring the evolutionary history of the plant NLR family, as previously suggested (Pan et al. 2000).

**Figure 2.**
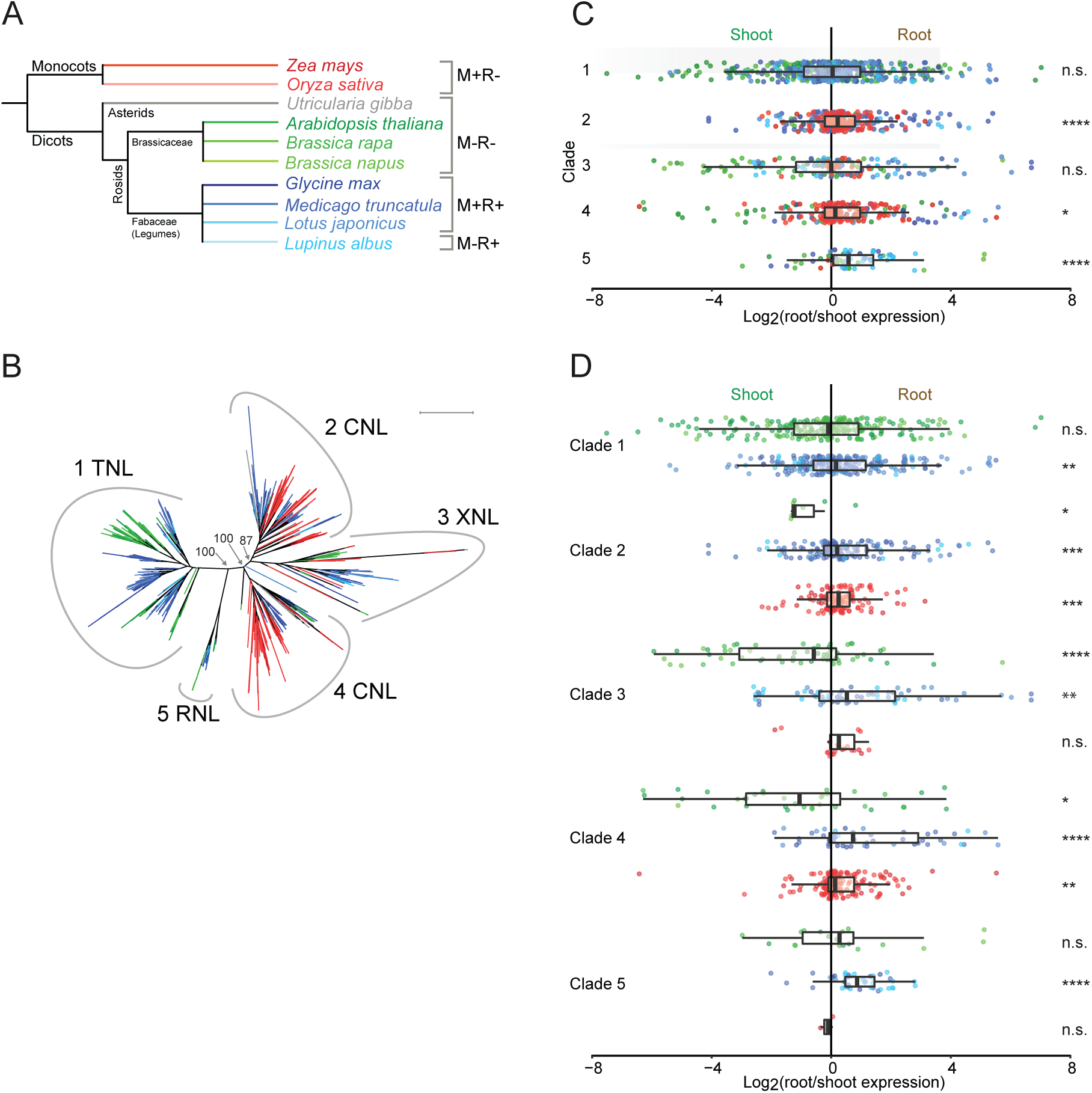
NBD protein expression patterns and sequence phylogeny. Colors indicate the plant species from which the NLR originates, with reference to **Figure 2A**. **A)** Species-level phylogenetic tree. The symbiotic status of each species is indicated on the right; M: Mycorrhiza. R: Rhizobia. +: engages in endosymbiosis. -: does not engage in endosymbiosis. **B)** Phylogenetic tree based on the NBD protein sequence of identified NLR genes in the species indicated in Figure 2A. Numbers at branches indicate bootstrap values for the branching of the 5 major clades. Peripheral numbers indicate clade designation, and NLR designation indicate enrichment of the corresponding NLR type in the given clade. Scale bar indicate 1.0 average amino acid substitutions per site. See **Supplemental File 2** for full bootstrap analysis of the tree. See Supplemental table 7 for NLR distribution at the clade and species level. **C)** Per clade log2 root/shoot expression ratios of the NLR genes shown in B) for which expression data is available. Each colored dot represents one NLR gene. Box plot bars show median with boxes indicating 25^th^ and 75^th^ percentiles and whiskers indicating 1.5 times the interquartile range. **D)** Same as C) but with expression data separated into groups depending on the species evolutionary descent colored according to Figure 2A. See Supplemental table 8 for *P*-values for inter-clade and inter-species differences.

In accordance with previous studies, we did not observe any sequences from monocots in the TNL-enriched clade 1, but among all sequences analysed we did find 7 monocot NLRs that had an identifiable TIR-like domain, which has previously been observed to be juxtaposed irregularly compared to the normal TIR domain (Meyers et al. 2002; Caplan et al. 2013). We did not recover any TIR or TIR-related domain containing NLR sequences from *U. gibba* either, despite it being a dicot (Pan et al. 2000; Fluhr 2001; Tarr et al. 2009; Ibarra-Laclette et al. 2013). In fact, *U. gibba* sequences were only found in clades 2 and 4 (Figure 2B and Supplemental table 7).

We then plotted NLR root/shoot ratios for the five NLR clades. Across data from all species, we observed highly significant root skews for the CNL-enriched clade 2 and for the RNL-enriched clade 5 (Figure 2C and Supplemental table 8). When examining the *Brassicaceae*, legume and monocot species groups separately, we found significant shoot skews for *Brassicaceae* clades 2, 3 and 4, and significant root skews for monocot clades 2 and 4 and for all legume NLR clades. The mean *Brassicaceae* root/shoot expression ratios deviated significantly from those of legumes for clades 1-4, and from monocots for clades 3 and 4 (Figure 2D and Supplemental table 8). In contrast, we did not find significant deviations between *Brassicaceae* and legumes for the RNL-enriched clade 5, where both species groups showed root-skewed expression. Since we observed significant *Brassicaceae* deviations for multiple NLR clades, a monophyletic group of NLRs was not responsible for the *Brassicaceae* expression shift. However, there were differences between the NLR clades in the severity of the shift, with the smallest effect seen for the TNL-enriched clade 1.

Comparing the species tree (Figure 2A) to the NBD-based NLR tree (Figure 2B), we noted that clade 1 and 5 in the NBD tree contained mainly dicot members, whereas clades 3 and 4 comprised monocot and dicot members from all species, in line with the species tree. In contrast, clade 2 from the NBD-tree was depleted in dicot *Brassicaceae* members, while both legume and monocot members were well-represented, indicating a family-specific depletion of a major NLR clade in the *Brassicaceae* family (Figure 2B and Supplemental table 7).

### NLR Clade 2 depletion is not generally associated with loss of mycorrhization

Although the NLR clade 2 depletion observed in the *Brassicaceae* family (Supplemental table 7) could not explain the *Brassicaceae* expression shift, it remained possible that NLR clade 2 would generally be depleted across non-mycorrhizal plants, pointing to a potentially specialized function in guarding the endomycorrhizal signalling machinery. To test this hypothesis, we identified and extracted NLR protein sequences from 8 additional non-mycorrhizal plant species and constructed a new phylogenetic tree containing a total of 2,448 NLR sequences (Figure 3A-B, Supplemental table 9, **and Supplemental file 3**). We found that 120 out of the 415 new NLR sequences were present in clade 2, leading us to reject our hypothesis that this clade had evolved specifically for guarding root endosymbiotic symbiotic components (Figure 3C and Supplemental table 9). After including three additional *Brassicaceae* species, we still observed a pronounced family-specific *Brassicaceae* depletion in clade 2, as we only found 20 out of 544 *Brassicaceae* NLRs belonging to this family (Supplemental table 9). In conclusion, NLR clade 2 depletion is likely *Brassicaceae* family specific and is not generally associated with loss of the endomycorrhizal pathway.

**Figure 3.**
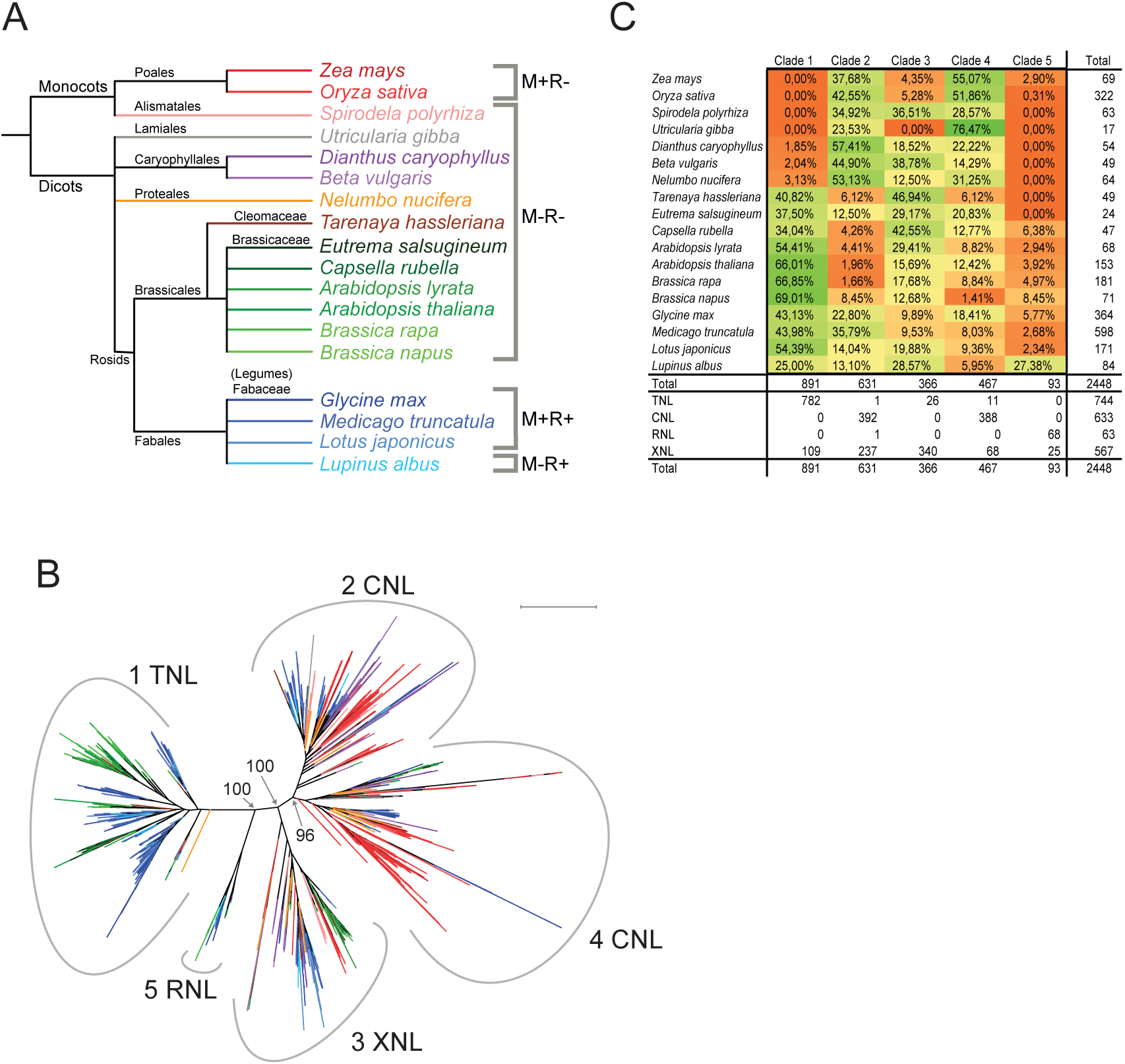
NBD phylogeny including additional non-mycorrhizal species. **A**) Species-level phylogenetic tree. The symbiotic status of each species is indicated on the right. M: Mycorrhiza. R: Rhizobia. +: engages in endosymbiosis. -: does not engage in endosymbiosis. **B**) Phylogenetic tree based on the NBD protein sequence of identified NLR genes in the species indicated in Figure 3A. Numbers at branches indicate bootstrap values for the branching of the 5 major clades. Peripheral numbers indicate clade designation. Scale bar indicate 1.0 average amino acid substitutions per site. Colors indicate the plant species from which the NBD originates, with reference to Figure 3A. See Supplemental File 3 for full bootstrap analysis of the tree. See Supplemental table 9 for NLR distribution at the clade and species level. **C**) Table showing the percentage of NLRs in each species, with respect to each clade shown in Figure 3B.

## Discussion

Cytoplasmic NLRs make up the last line of defence against potentially pathogenic microbes that have evaded physical barriers and membrane-localized PRRs to successfully deliver effectors into plant cells. The stable root/shoot NLR expression ratios observed here are consistent with a defence system in which NLR expression patterns are hardwired to match organ-specific effector challenges, in anticipation of microbial challenge, similar to that observed for the plant circadian cycle (Ingle 2011; Wang et al. 2011). Indeed, we also found that rhizobium inoculation of the nodulating legume *Lotus* did not alter the overall pattern of NLR expression, further underlining the stability within species of NLR root/shoot expression ratios. It was striking that we found an overall root-skew in NLR expression in the majority of plant species. This suggested that roots generally experience a higher level of effector pressure than shoots, despite the fact that NLR function has mainly been characterized in the context of shoot-pathogen interactions (Erb et al. 2009; Nishimura and Dangl 2010). It might not be surprising given the complexity of soil microbial communities, but our data does underline the need for establishing new root pathosystems and for understanding the role of NLRs in root-microbe interactions.

Plants from the *Brassicaceae* family made up a very conspicuous group of outliers that displayed shoot-rather than root-skewed NLR expression. The *Brassicaceae* are also outliers in the sense that they have lost the capacity for root endomycorrhization, which remains functional in 80-90% of land plants (Parniske 2008; Delaux et al. 2014). This symbiotic interaction between plant roots and arbuscular mycorrhizal fungi has existed for around 400 million years, coinciding with the appearance of terrestrial plants, and parts of the mycorrhization signalling machinery have been recruited in the ∼110 million year old symbiotic interaction between plants and nitrogen fixing rhizobia (Parniske 2000; Deguchi et al. 2007). NLRs are also found in early land plant species, such as Bryophytes and lycophytes (Xue et al. 2012; Yue et al. 2012; Jacob et al. 2013; Tanigaki et al. 2014), meaning that endomycorrhizal signalling has co-evolved with NLRs through hundreds of millions of years.

It is conceivable that a specialized set of phylogenetically related NLRs could have evolved specifically to guard the root endosymbiotic machinery or other root specific pathways, and that the *Brassicaceae* NLR expression shift might be caused by the loss of such a group of NLRs. Here, we tested this hypothesis by grouping NLRs according to the sequence homology of their NBD domains, identifying five major clades. While the CNL-enriched NLR clade 2 was strongly depleted in the *Brassicaceae*, it was well-represented in other non-mycorrhizal plants. In addition, we observed a *Brassicaceae* shoot skew for all NLR clades, with the smallest shift observed for TNLs, which are absent in the endomycorrhizal monocots rice and maize, and therefore cannot be generally required for protecting the endomycorrhizal signalling machinery. The shift in *Brassicacae* NLR expression could thus not be attributed to the loss of a single NLR clade, and our data did not support the existence of a specific group of phylogenetically distinct NLRs guarding the root endosymbiotic machinery.

The general expression shift towards the shoot across four major NLR clades suggests a reduced anticipation of effector challenge to root cells relative to shoot cells in the *Brassicaceae*. We envisage two scenarios, which are not mutually exclusive, that could account for the shift. First, our data is consistent with a model where NLRs were randomly recruited from an expanding NLR complement, regardless of phylogenetic origin, for guarding root specific components. When the guarded pathways became defunct in the *Brassicaceae* family, it gradually lost the associated root-expressed NLRs across the different NLR clades, leading to the overall shoot skew in NLR expression. Second, rather than passively reducing the effector challenge level to roots by loss of a potentially exposed pathway, the *Brassicaceae* could have developed family-specific active measures that efficiently deter putative soil pathogens before they have a chance to deploy their effectors, reducing the requirement for NLR protection. One possibility is that the *Brassicaceae* maintain high levels of antimicrobial glucosinolates in the root apoplasm, and there are indications that root have higher constitutive glucosinolate levels than shoots (Van Dam, Tytgat, and Kirkegaard 2009). Another is that the *Brassicaceae* have evolved a unique set of highly efficient pattern recognition receptors that quickly eliminate putative root pathogens. For instance, the Ef-Tu and lipopolysaccharide PRRs are thought to be *Brassicaceae*-specific (Kunze et al. 2004; Ranf et al. 2015).

The root endosymbiosis signalling pathway allows intracellular accommodation of symbiotic mycorrhizal fungi and rhizobia (Madsen et al. 2010; Oldroyd 2013). This could impose severe constraints on the general defence mechanisms employed in roots of plant species that rely on symbiotic interactions for nutrient acquisition, compelling these symbiotic species to depend to a greater extent on NLR effector recognition in roots. We propose that the loss of root endomycorrhizal signalling in the *Brassicaceae* family offers the most parsimonious explanation for the *Brassicaceae* NLR expression shift. Its loss would both have removed a potentially heavily NLR-guarded pathway and eliminated constraints impeding development of more effective general root defence systems. This hypothesis is consistent with both scenarios described above, agrees with the discovery of apparently *Brassicaceae*-specific PRRs (Kunze et al. 2004; Ranf et al. 2015), and suggests that *Brassicaceae*, and perhaps other non-mycorrhizal plants, may be rich sources of unique PRRs and antimicrobial root metabolites.

## Materials and methods

### Identification of putative NLR genes

To allow identification of putative NLR genes, protein sequences were downloaded as indicated (Supplemental table 1). Annotation versions were chosen for compatibility with the available microarray or RNA-seq data to allow subsequent expression analysis. This is why the latest versions were not used in all cases. NLR genes were then identified in a three-step procedure. First, candidate genes were selected using HMMER 3.1b1 (Eddy 2011) based on the NB-ARC PFAM protein domain PF00931. Second, the candidate list was filtered by performing a search for conserved protein domains using CDD (Marchler-Bauer et al. 2011), requiring that the selected putative NLR genes contain, in addition to the NB-ARC domain, either LRR, TIR, PLN00113, PLN03194, or PLN03210 domains. Third, all NLR gene sequences were manually curated to identify and remove false positives. The total number of identified NLR genes in each of the 18 species is shown in Supplemental table 3, with sequences available in **Supplemental file 4**.

### *Lotus* RNA-seq

*L. japonicus* ecotype Gifu (Handberg and Stougaard 1992) seeds were surface sterilized, germinated and grown in conditions as described previously (Kawaharada et al. 2015). Three biological replicates per sample were analyzed with each consisting of 10 seedlings grown on 1/4 B&D plates for 10 days before inoculation of the roots with 750 μL of an *M. loti* R7A suspension (OD_600_ = 0.02) or water. Three days post-inoculation roots and shoots were separated and total RNA was isolated using a NucleoSpin^®^ RNA Plant kit (Machery-Nagel) according to the manufacturer’s instructions. RNA quality was assessed with on an Agilent 2100 Bioanalyser and samples were sent to GATC Biotech (http://gatc-biotech.com/) for library preparation and sequencing. Sequencing data have been deposited at the NCBI Short Read Archive with BioProject ID PRJNA384655 and are available for analysis on *Lotus* Base (Mun et al. 2016).

### Analysis of NLR gene expression data

For tissues-specific gene expression enrichment analysis (Figure 1 A-D), we classified genes as being enriched in a specific tissue group, if the average expression level in a that group was higher than the average of all other tissue groups, and at least two times higher than that of at least one other tissue group.

In order to evaluate root/shoot expression ratios, available expression data was downloaded as indicated in Supplemental table 2. Samples IDs along with expression values are available in **Supplemental file 1**. For *Lotus* and *B. rapa*, probes were reassigned to the updated annotation using BLAST to match probe and cDNA sequences (e-value cut-off 0.001), assigning only the best matching probe to a gene. For *Lotus*, *Medicago* and soybean, samples representing identical or closely related plant accessions were used in the analysis. For rice and maize, data from a number of different accessions were used, but only data where both root and shoot samples had been assayed within the same experiment were used to ensure the comparability of samples from the two tissues. For *B. napus*, the analysis was based on raw RNA-seq reads. RNA-seq data files were downloaded from the NCBI short read archive (https://www.ncbi.nlm.nih.gov/sra) and reads from each library were assembled using Trinity (--full_cleanup) (Haas et al. 2013) followed by clustering using cd-hit-est v.4.6.6 (-M 16000-T 8) (Fu et al. 2012). Next, the longest open reading frames were identified for each transcript and the corresponding protein sequences were used for identification of NLRs as described. Reads were mapped back to the gene set output from cd-hit-est using STAR (--runMode genomeGenerate --genomeChrBinNbits 14) parameters for index generation and standard options for mapping (Dobin et al. 2013). Finally reads mapping to multiple locations were filtered out followed by summarizing read counts per gene for each sample. For all species, expression data from the genes with the 15% lowest expression levels were filtered out, and the log2 NLR root/shoot expression ratios were normalized by subtracting the mean value for all genes. Expression ratios were plotted using ggplot2 in R version 3.1.2.

The significance of differences in mean expression ratios between all genes and NLR genes were evaluated using Student’s t-test (Supplemental table 4). Next, the significances of interspecies differences in root/shoot expression ratios were evaluated using one-way ANOVA followed by Tukey’s multiple comparison test as implemented in GraphPad Prism 6 (Supplemental table 5). Differences in the average root/shoot expression by NLR gene clade or domain based on the phylogenetic tree shown in Figure 2B, were evaluated using one-way ANOVA followed by Tukey’s multiple comparison test, or Student’s t-test, as implemented in GraphPad Prism 6 (Supplemental tables 6 and 8).

### Construction of NLR protein phylogeny

Sequences of the NB-ARC domains of identified R genes were extracted using a python script, based on domains as identified by the CCD search, and aligned using Clustal Omega v1.2.3 (Sievers et al. 2011). Sequences were then filtered for low coverage positions (50% cut-off) and sequences lacking more than 50% of the aligned NB-ARC domain were removed. Phylogenetic trees were constructed in IQ-Tree v.1.5.2 and evaluated using the ultrafast bootstrap approximation approach (UFBoot) implemented the software package (Minh et al. 2013; L.-T. Nguyen et al. 2015). The resulting tree was colored by species using colorTree v1.1 and visualized using Dendroscope v3.5.7 ( Chen et al. 2009; Huson et al. 2012). See **Supplemental files 2 and 3** for NB-ARC domain alignments of the trees described in Figures 2B and 3B respectively, along with bootstrap analysis. See **Supplemental file 4** for sequences for all NLRs used to construct the phylogenetic trees, and **Supplemental file 5** for a general overview of all NLRs used in the study.

## Author contributions

DM, VG, AB, TM, WB, and SUA analysed data. SK carried out the *Lotus* RNA-seq experiment. SUA designed and supervised the study. DM and SUA wrote the manuscript.

## Acknowledgements

This work was supported by the Danish National Research Foundation grant no. DNRF79. The authors wish to acknowledge all research groups contributing expression data used in our meta-analysis.

## Supplemental Information

### Supplemental figures

**Supplemental figure 1.** Density plots displaying the distribution of the normalized logarithm of the root/shoot expression ratios of all genes and NLR genes in *Lotus*, from two different data sets, excluding or including inoculated samples.

**Supplemental figure 2.** Density plots displaying the distribution of the normalized logarithm of the root/shoot expression ratios of all genes and NLR genes for the species and experiments indicated.

**Supplemental figure 3.** Log root/shoot expression ratios split by NLR domain type.

### Supplemental files

**Supplemental file 1.** Complete set of expression data for all species.

**Supplemental file 2**. NB-ARC alignment of sequences used to construct the phylogenetic tree in Figure 2B, along with bootstrap analysis and the resulting phylogenetic tree.

**Supplemental file 3**. NB-ARC alignment of sequences used to construct the phylogenetic tree in Figure 3B, along with bootstrap analysis and the resulting phylogenetic tree.

**Supplemental file 4.** Full length sequences for all NLRs identified and used in this study. **Supplemental file 5.** Full list of NLR genes identified, with domains, designations and normalized log_2_ root/shoot expression ratios.

### Supplemental tables

**Supplemental table 1**. Sources of the protein sequences used in the NLR analysis.

**Supplemental table 2**. Expression data sources used in the NLR analysis. **Supplemental table 3**. Number of NLR genes identified through computational analyses for all species where expression analysis was carried out.

**Supplemental table 4.** Mean root/shoot gene expression ratios for all genes and NLR genes.

**Supplemental table 5**. Cross-species comparison of normalized log_2_ root/shoot NLR expression ratios supporting Figure 1I. ANOVA and Tukey’s multiple comparison test was used for calculation of *P*-values.

**Supplemental table 6.** Cross-species domain comparison of normalized log_2_ root/shoot NLR gene expression ratios for Supplemental figure 3. ANOVA and Tukey’s multiple comparison test was used for calculation of *P*-values.

**Supplemental table 7**. Clade distribution of NLR genes for the phylogenetic tree used in Figure 2B, including number of identified NLRs identified with at least one TIR (TNL), CC (CNL) or CC_R_ (RNL) domain, or none of the former three (XNL), and their clade distribution patterns, with reference to Figure 2B.

**Supplemental table 8.** Cross-species clade comparison of normalized log_2_ root/shoot NLR gene expression ratios for Figure 2C and 2D. ANOVA and Tukey’s multiple comparison test was used for calculation of *P*-values.

**Supplemental table 9**. Clade distribution of NLR genes for the phylogenetic trees used in Figure 3B, including number of identified NLRs identified with at least one TIR (TNL), CC (CNL) or CC_R_ (RNL) domain, or none of the former three (XNL), and their clade distribution patterns, with reference to Figure 3B.

## Supplemental figure

**Supplemental Figure 1.**
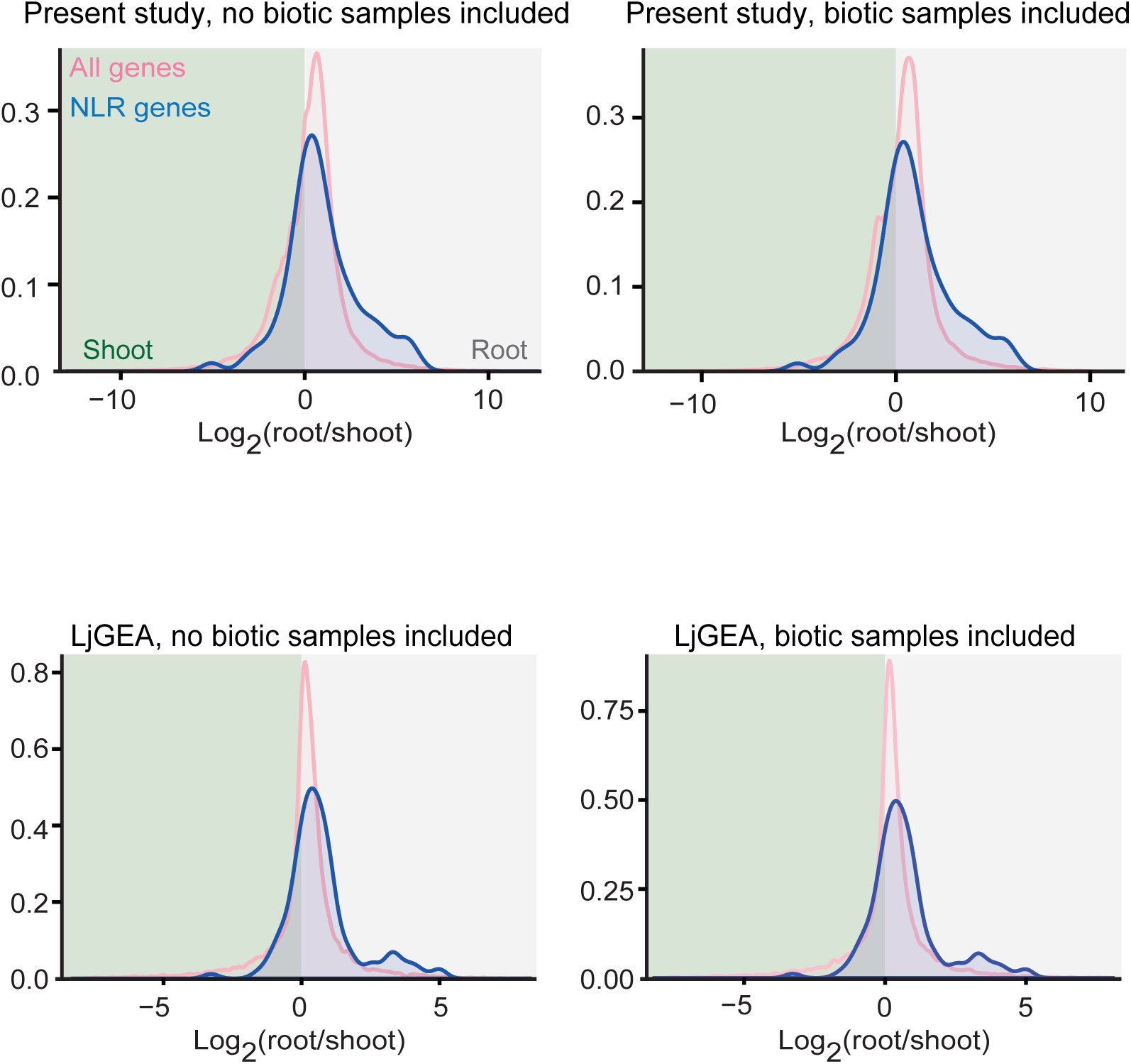
Density plots displaying the distribution of the logarithm of the root/shoot expression ratios of all genes and NLR genes in Lotus, from two different data sets, excluding or including inoculated samples.

**Supplemental figure 2.**
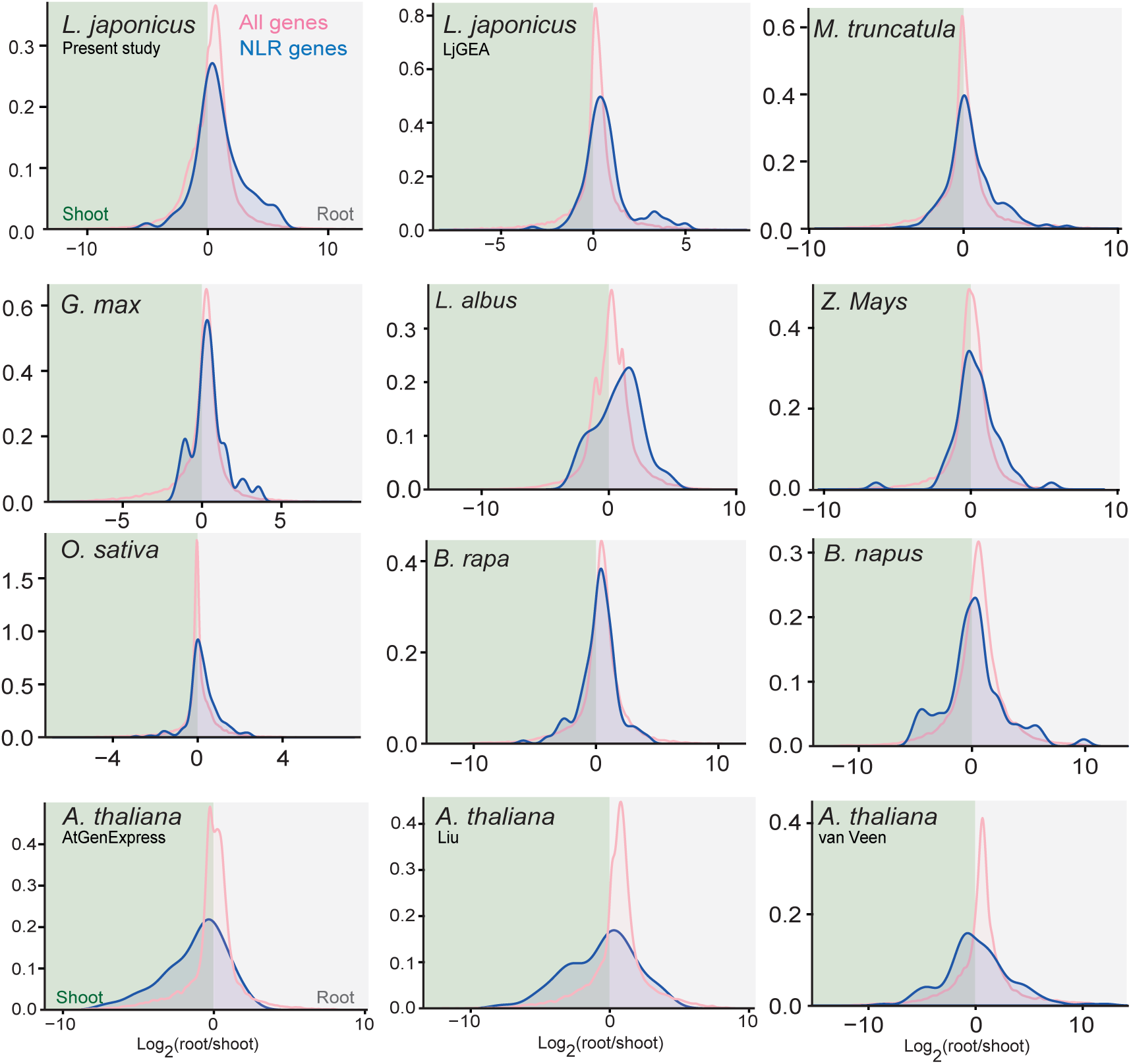
Density plots displaying the distribution of the logarithm of the root/shoot expression ratios of all genes and NLR genes for the species indicated.

**Supplemental figure 3.**
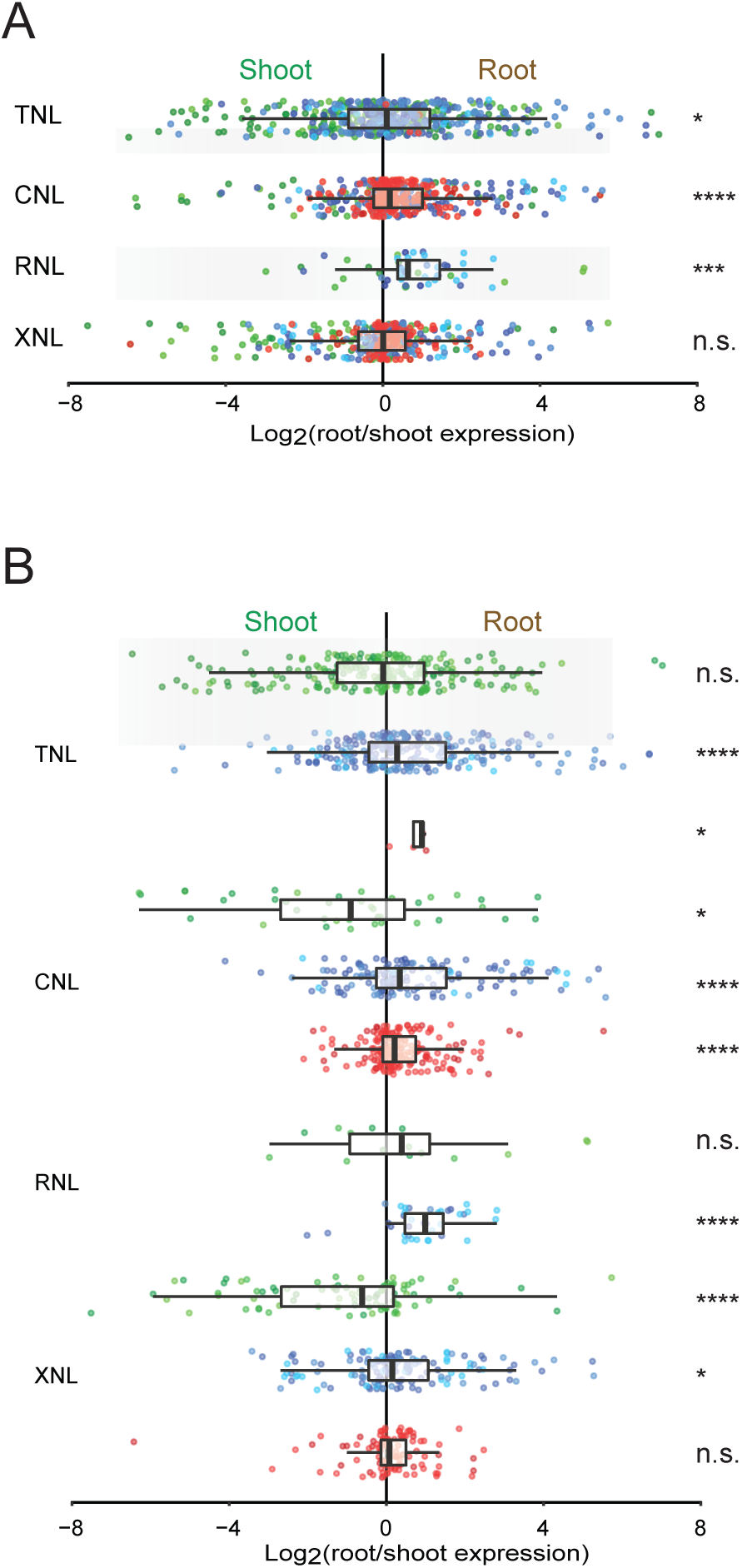
Colors indicate the plant species from which the NLR originates, with reference to Figure 2A. **A)** Log2 root/shoot expression ratios. Each colored dot represents one NLR gene. Box plot bars show median, with boxes indicating 25^th^ and 75^th^ percentiles and whiskers indicating 1.5 times the interquartile range. **C)** Same as B) but with expression data separated into groups depending on the species evolutionary descent colored according to Figure 2A.

## Supplemental tables

**Supplemental table 1.** Sources of the protein sequences used in the NLR analyses. *: The *L. albus* analysis was based on *de novo* assembled transcripts and not on annotated protein coding genes.

| Species | Version | Source |
| --- | --- | --- |
| *A. thaliana* | 10 | ftp://ftp.arabidopsis.org/home/tair/Sequences/blast_datasets/TAIR10_blastsets/TAIR10_pep_20101214_updated |
| *B. napus* | - | http://www.ncbi.nlm.nih.gov/sra/SRP028575 |
| *B. rapa* | 1.2 | http://www.plantgdb.org/download/Download/xGDB/BrGDB/Brapa_197_peptide.fa.gz |
| *G. max* | 1.09 | http://www.plantgdb.org/download/Download/xGDB/GmGDB/Gmax_109_peptide.fa.gz |
| *L. japonicus* | 3.0 | http://www.kazusa.or.jp/lotus/ |
| *L. albus* | * | http://lupal.comparative-legumes.org/data/2013/797f6de2ea8102be4b875c165ab5e994/LAGI01_express_annotate.txt |
| *M. truncatula* | 3.5 | ftp://ftp.jcvi.org/pub/data/m_truncatula/Mt3.5/Annotation/Mt3.5v5/Mt3.5v5_GenesProteinSeq_20111014.fa |
| *O. sativa* | 7.0 | ftp://ftp.plantbiology.msu.edu/pub/data/Eukaryotic_Projects/o_sativa/annotation_dbs/pseudomolecules/version_7.0/all.dir/all.pep |
| *Z. mays* | 3 | http://plantgdb.org/download/download.php?dir=/PublicPlantSeq/Dump/Z/Zea_mays |
| *U. gibba* | 4.1 | http://de.iplantcollaborative.org/dl/d/4E6FBB33-2FE7-4404-8548-D3BC14E2CEFC/Utricularia_gibba-ft-CDS-gid-19475-prot.fasta |

**Supplemental table 2.**
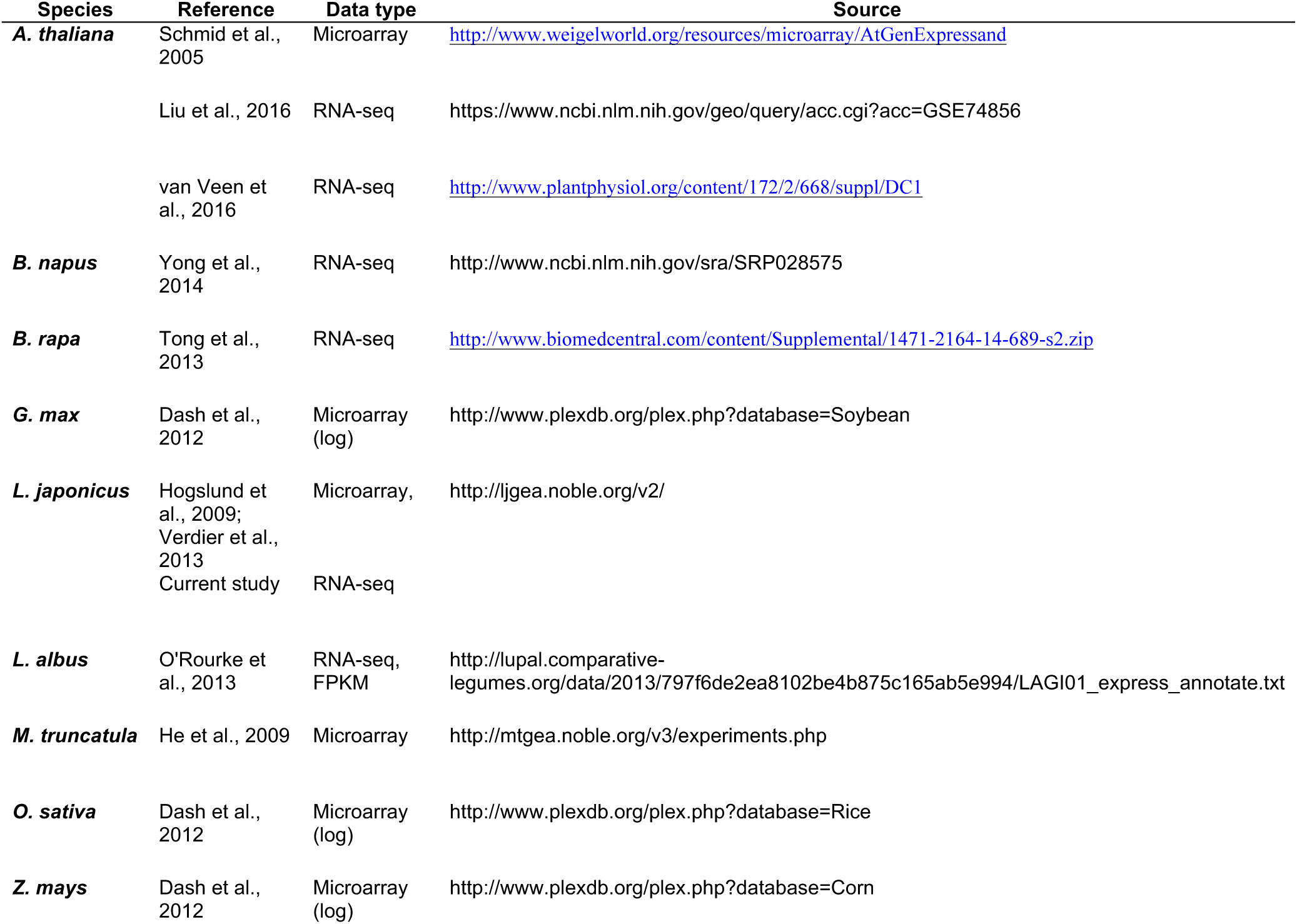
Expression data sources used in the NLR gene analysis. Please refer to the data sources listed for a detailed description of the samples.

**Supplemental table 3.**
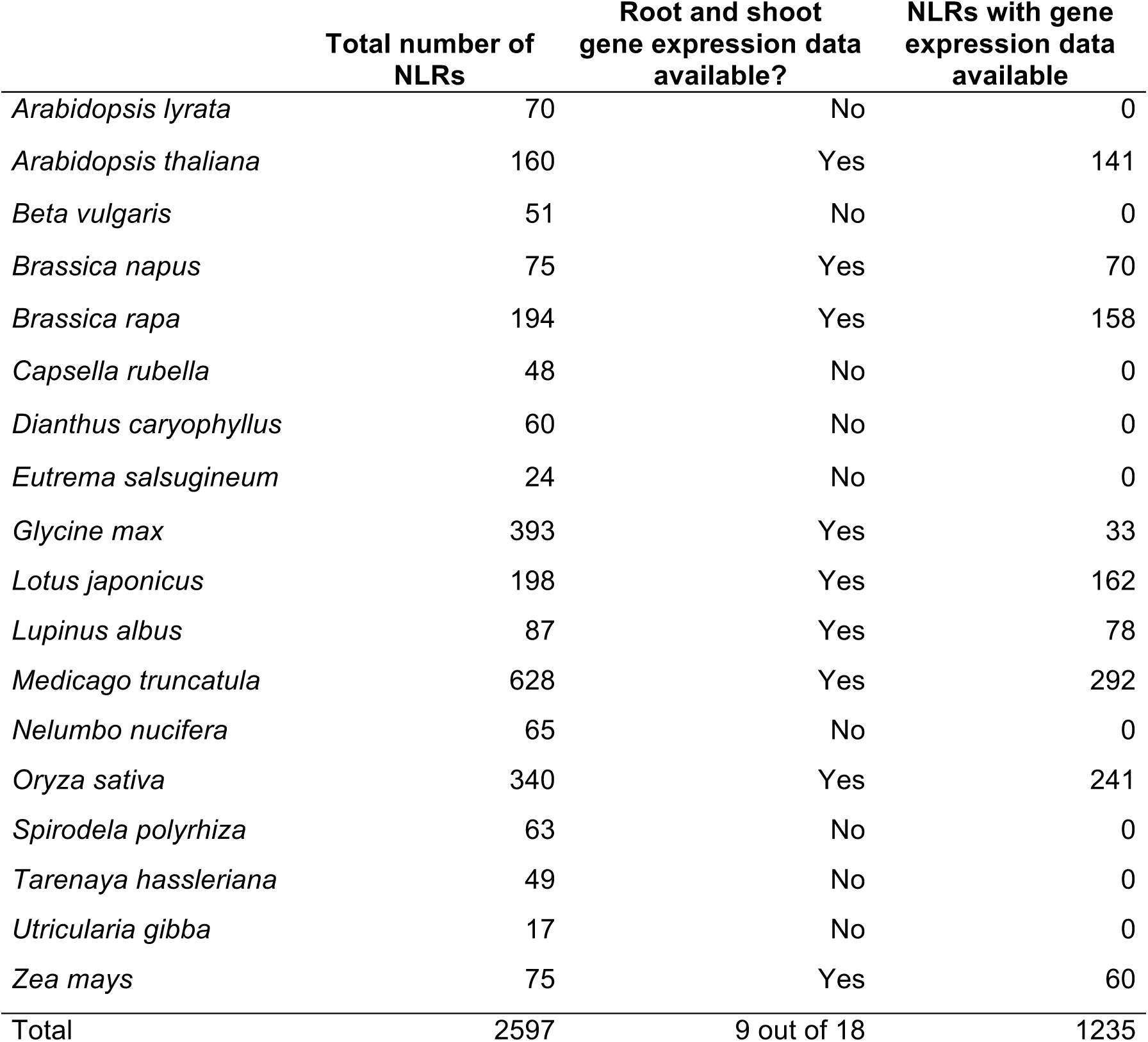
Number of NLR genes identified through computational analyses for all species. Expression analysis was carried out where root and shoot expression data was available for the indicated number of NLRs with expression data available. The numbers in this table include sequences that were subsequently filtered out as explained in the Materials and Methods section.

**Supplemental Table 4.**
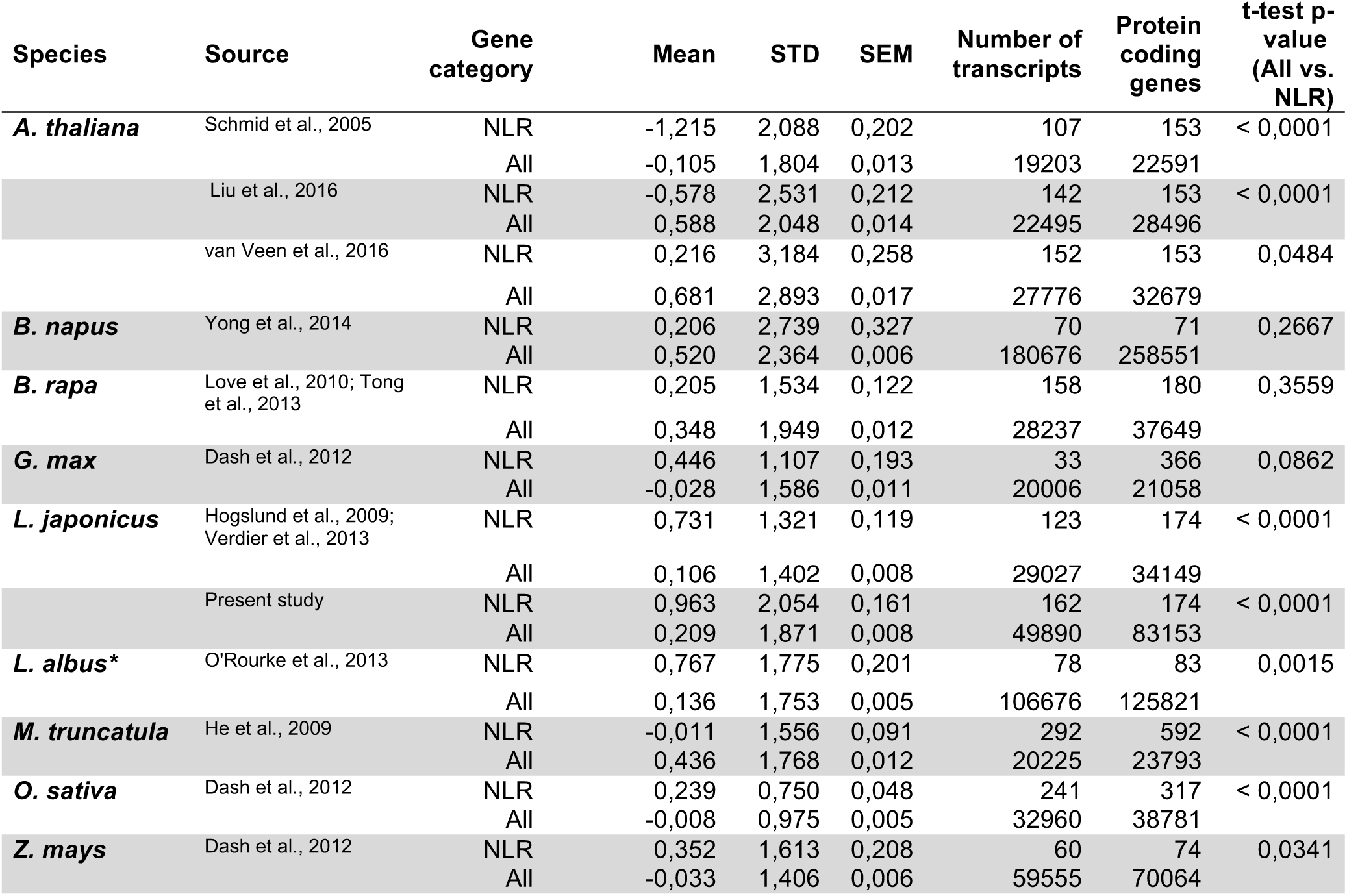
Root/shoot mean gene expression ratios for NLR and all genes. Mean: average log_2_(root/shoot expression). STD: standard deviation. SEM: Standard error of the mean. N: Number of genes with expression values included in the analysis. *: The *L. albus* analysis was based on *de novo* assembled transcripts and not on annotated protein coding genes.

**Supplemental Table 5.** Cross-species comparison of root/shoot NLR expression ratios for Figure 1I. ANOVA and Tukey’s multiple comparison test was used for calculation of *P*-values. ns: *P* > 0.05; *: *P* ≤ 0.05; **: *P* ≤ 0.01; ***: *P* ≤ 0.001; ****: *P* ≤ 0.0001.

| **Tukey's multiple comparisons test** | **Mean Diff.** | **95% CI of diff.** | **Significant** | **Summary** | **Adjusted P Value** |
| --- | --- | --- | --- | --- | --- |
| *Lotus, present study vs. Lotus, LjGEA* | 0,1203 | -0,6249 to 0,8656 | No | ns | > 0,9999 |
| *Lotus, present study vs. Soybean* | 0,2713 | -0,9188 to 1,461 | No | ns | 0,9999 |
| *Lotus, present study vs. Medicago* | 0,2834 | -0,3271 to 0,8939 | No | ns | 0,9355 |
| *Lotus, present study vs. Lupin* | 0,1142 | -0,7446 to 0,9731 | No | ns | > 0,9999 |
| *Lotus, present study vs. Arabidopsis, van Veen* | 1,21 | 0,5062 to 1,914 | Yes | **** | < 0,0001 |
| *Lotus, present study vs. Arabidopsis, Liu* | 1,911 | 1,195 to 2,628 | Yes | **** | < 0,0001 |
| *Lotus, present study vs. Arabidopsis, AtGenExpress* | 1,855 | 1,079 to 2,632 | Yes | **** | < 0,0001 |
| *Lotus, present study vs. B. rapa* | 0,8887 | 0,1919 to 1,585 | Yes | ** | 0,0019 |
| *Lotus, present study vs. B. napus* | 1,059 | 0,1679 to 1,951 | Yes | ** | 0,0059 |
| *Lotus, present study vs. Rice* | 0,4983 | -0,1348 to 1,131 | No | ns | 0,2942 |
| *Lotus, present study vs. Maize* | 0,3604 | -0,5814 to 1,302 | No | ns | 0,9845 |
| *Lotus, LjGEA vs. Soybean* | 0,151 | -1,071 to 1,373 | No | ns | > 0,9999 |
| *Lotus, LjGEA vs. Medicago* | 0,163 | -0,5068 to 0,8329 | No | ns | 0,9997 |
| *Lotus, LjGEA vs. Lupin* | -0,006107 | -0,9081 to 0,8959 | No | ns | > 0,9999 |
| *Lotus, LjGEA vs. Arabidopsis, van Veen* | 1,09 | 0,3338 to 1,845 | Yes | *** | 0,0002 |
| *Lotus, LjGEA vs. Arabidopsis, Liu* | 1,791 | 1,023 to 2,559 | Yes | **** | < 0,0001 |
| *Lotus, LjGEA vs. Arabidopsis, AtGenExpress* | 1,735 | 0,9112 to 2,559 | Yes | **** | < 0,0001 |
| *Lotus, LjGEA vs. B. rapa* | 0,7683 | 0,01901 to 1,518 | Yes | * | 0,0386 |
| *Lotus, LjGEA vs. B. napus* | 0,9389 | 0,005914 to 1,872 | Yes | * | 0,0469 |
| *Lotus, LjGEA vs. Rice* | 0,378 | -0,3126 to 1,069 | No | ns | 0,8228 |
| *Lotus, LjGEA vs. Maize* | 0,2401 | -0,7412 to 1,221 | No | ns | 0,9997 |
| *Soybean vs. Medicago* | 0,01204 | -1,132 to 1,156 | No | ns | > 0,9999 |
| *Soybean vs. Lupin* | -0,1571 | -1,451 to 1,137 | No | ns | > 0,9999 |
| *Soybean vs. Arabidopsis, van Veen* | 0,9386 | -0,2582 to 2,135 | No | ns | 0,2996 |
| *Soybean vs. Arabidopsis, Liu* | 1,64 | 0,4357 to 2,844 | Yes | *** | 0,0005 |
| *Soybean vs. Arabidopsis, AtGenExpress* | 1,584 | 0,3432 to 2,825 | Yes | ** | 0,0018 |
| *Soybean vs. B. rapa* | 0,6174 | -0,5754 to 1,810 | No | ns | 0,8711 |
| *Soybean vs. B. napus* | 0,7879 | -0,5280 to 2,104 | No | ns | 0,7208 |
| *Soybean vs. Rice* | 0,227 | -0,9297 to 1,384 | No | ns | > 0,9999 |
| *Soybean vs. Maize* | 0,08906 | -1,262 to 1,440 | No | ns | > 0,9999 |
| *Medicago vs. Lupin* | -0,1691 | -0,9634 to 0,6251 | No | ns | > 0,9999 |
| *Medicago vs. Arabidopsis, van Veen* | 0,9265 | 0,3033 to 1,550 | Yes | **** | < 0,0001 |
| *Medicago vs. Arabidopsis, Liu* | 1,628 | 0,9904 to 2,266 | Yes | **** | < 0,0001 |
| *Medicago vs. Arabidopsis, AtGenExpress* | 1,572 | 0,8677 to 2,276 | Yes | **** | < 0,0001 |
| *Medicago* vs. *B. rapa* | 0,6053 | -0,01014 to 1,221 | No | ns | 0,0589 |
| *Medicago* vs. *B. napus* | 0,7759 | -0,05344 to 1,605 | No | ns | 0,0924 |
| *Medicago* vs. *Rice* | 0,215 | -0,3274 to 0,7573 | No | ns | 0,9795 |
| *Medicago* vs. *Maize* | 0,07702 | -0,8063 to 0,9603 | No | ns | > 0,9999 |
| *Lupin* vs. *Arabidopsis*, van Veen | 1,096 | 0,2277 to 1,964 | Yes | ** | 0,0022 |
| *Lupin* vs. *Arabidopsis*, Liu | 1,797 | 0,9188 to 2,675 | Yes | **** | < 0,0001 |
| *Lupin* vs. *Arabidopsis*, AtGenExpress | 1,741 | 0,8133 to 2,669 | Yes | **** | < 0,0001 |
| *Lupin* vs. *B. rapa* | 0,7745 | -0,08790 to 1,637 | No | ns | 0,1282 |
| *Lupin* vs. *B. napus* | 0,945 | -0,08096 to 1,971 | No | ns | 0,1052 |
| *Lupin* vs. *Rice* | 0,3841 | -0,4277 to 1,196 | No | ns | 0,9266 |
| *Lupin* vs. *Maize* | 0,2462 | -0,8239 to 1,316 | No | ns | 0,9998 |
| *Arabidopsis*, van Veen vs. *Arabidopsis*, Liu | 0,7014 | -0,02587 to 1,429 | No | ns | 0,0708 |
| *Arabidopsis*, van Veen vs. *Arabidopsis*, AtGenExpress | 0,6454 | -0,1410 to 1,432 | No | ns | 0,234 |
| *Arabidopsis*, van Veen vs. *B. rapa* | -0,3212 | -1,029 to 0,3868 | No | ns | 0,9448 |
| *Arabidopsis*, van Veen vs. *B. napus* | -0,1507 | -1,051 to 0,7495 | No | ns | > 0,9999 |
| *Arabidopsis*, van Veen vs. *Rice* | -0,7116 | -1,357 to -0,06612 | Yes | * | 0,0166 |
| *Arabidopsis*, van Veen vs. *Maize* | -0,8495 | -1,800 to 0,1006 | No | ns | 0,1326 |
| *Arabidopsis*, Liu vs. *Arabidopsis*, AtGenExpress | -0,056 | -0,8538 to 0,7418 | No | ns | > 0,9999 |
| *Arabidopsis*, Liu vs. *B. rapa* | -1,023 | -1,743 to -0,3021 | Yes | *** | 0,0002 |
| *Arabidopsis*, Liu vs. *B. napus* | -0,8521 | -1,762 to 0,05800 | No | ns | 0,0918 |
| *Arabidopsis*, Liu vs. *Rice* | -1,413 | -2,072 to -0,7538 | Yes | **** | < 0,0001 |
| *Arabidopsis*, Liu vs. *Maize* | -1,551 | -2,510 to -0,5914 | Yes | **** | < 0,0001 |
| *Arabidopsis*, AtGenExpress vs. *B. rapa* | -0,9667 | -1,747 to -0,1864 | Yes | ** | 0,0031 |
| *Arabidopsis*, AtGenExpress vs. *B. napus* | -0,7961 | -1,754 to 0,1619 | No | ns | 0,2173 |
| *Arabidopsis*, AtGenExpress vs. *Rice* | -1,357 | -2,081 to -0,6331 | Yes | **** | < 0,0001 |
| *Arabidopsis*, AtGenExpress vs. *Maize* | -1,495 | -2,500 to -0,4899 | Yes | **** | < 0,0001 |
| *B. rapa* vs. *B. napus* | 0,1706 | -0,7242 to 1,065 | No | ns | > 0,9999 |
| *B. rapa* vs. *Rice* | -0,3904 | -1,028 to 0,2476 | No | ns | 0,6917 |
| *B. rapa* vs. *Maize* | -0,5283 | -1,473 to 0,4167 | No | ns | 0,8016 |
| *B. napus* vs. *Rice* | -0,5609 | -1,407 to 0,2852 | No | ns | 0,5721 |
| *B. napus* vs. *Maize* | -0,6989 | -1,795 to 0,3975 | No | ns | 0,6328 |
| *Rice* vs. *Maize* | -0,1379 | -1,037 to 0,7612 | No | ns | > 0,9999 |

**Supplemental Table 5.** Cross-species comparison of root/shoot NLR expression ratios for Figure 1I. ANOVA and Tukey's multiple comparison test was used for calculation of *P*-values. ns: $P > 0.05$ ; \*: $P \leq 0.05$ ; \*\*: $P \leq 0.01$ ; \*\*\*: $P \leq 0.001$ ; \*\*\*\*: $P \leq 0.0001$ .
| Tukey's multiple comparisons test | Mean Diff, | 95% CI of diff, | Significant | Summary |
| --- | --- | --- | --- | --- |
| *Brassicaceae* vs. *Legumes* | -1,207 | -1,458 to -0,9556 | Yes | **** |
| *Brassicaceae* vs. *Monocots* | -0,9098 | -1,229 to -0,5905 | Yes | **** |
| *Legumes* vs. *Monocots* | 0,2969 | -0,01776 to 0,6115 | No | ns |

**Supplemental table 6.** Cross-species domain comparison of normalized log_2_ root/shoot NLR gene expression ratios, for Supplemental figure 3. ANOVA and Tukey’s multiple comparison test was used for calculation of *P*-values.

|  | **P value (two tailed)** | **Significant (p=0.05)** |
| --- | --- | --- |
| All NLRs | 0,0004 | ** |
| All TNL | 0,0343 | * |
| All CNL | < 0,0001 | **** |
| All RNL | 0,0002 | *** |
| All XNL | 0,1902 | ns |
| Brassicaceae TNL | 0,1951 | ns |
| Brassicaceae CNL | 0,0139 | * |
| Brassicaceae RNL | 0,3267 | ns |
| Brassicaceae XNL | < 0,0001 | **** |
| Legume TNL | < 0,0001 | **** |
| Legume CNL | < 0,0001 | **** |
| Legume RNL | < 0,0001 | **** |
| Legume XNL | 0,0286 | * |
| Monocot TNL | 0,0131 | * |
| Monocot CNL | < 0,0001 | **** |
| Monocot XNL | 0,3712 | ns |

| **Tukey's multiple comparisons test** | **Mean Diff.** | **95% CI of diff.** | **Summary** |
| --- | --- | --- | --- |
| Brassicaceae TNL vs. Legumes TNL | -0,7424 | -1.181 to -0.3038 | *** |
| Brassicaceae TNL vs. Monocots TNL | -0,9144 | -3.076 to 1.247 | ns |
| Legumes TNL vs. Monocots TNL | -0,1719 | -2.331 to 1.987 | ns |
| Brassicaceae CNL vs. Legumes CNL | -1,798 | -2.447 to -1.148 | **** |
| Brassicaceae CNL vs. Monocots CNL | -1,442 | -2.075 to -0.8088 | **** |
| Legumes CNL vs. Monocots CNL | 0,3556 | -0.04291 to 0.7541 | ns |

| **Unpaired t-test** | **t, df** | **One- or two-tailed?** | **P value** |
| --- | --- | --- | --- |
| Legumes RNL vs. Brassicaceae RNL | t=0.8081 df=50 | Two-tailed | ns |

| **Tukey's multiple comparisons test** | **Mean Diff.** | **95% CI of diff.** | **Summary** |
| --- | --- | --- | --- |
| Brassicaceae XNL vs. Legumes XNL | -1,39 | -1.921 to -0.8596 | **** |
| Brassicaceae XNL vs. Monocots XNL | -1,18 | -1.740 to -0.6198 | **** |
| Legumes XNL vs. Monocots XNL | 0,2107 | -0.2855 to 0.7070 | ns |

**Supplemental Table 7.**
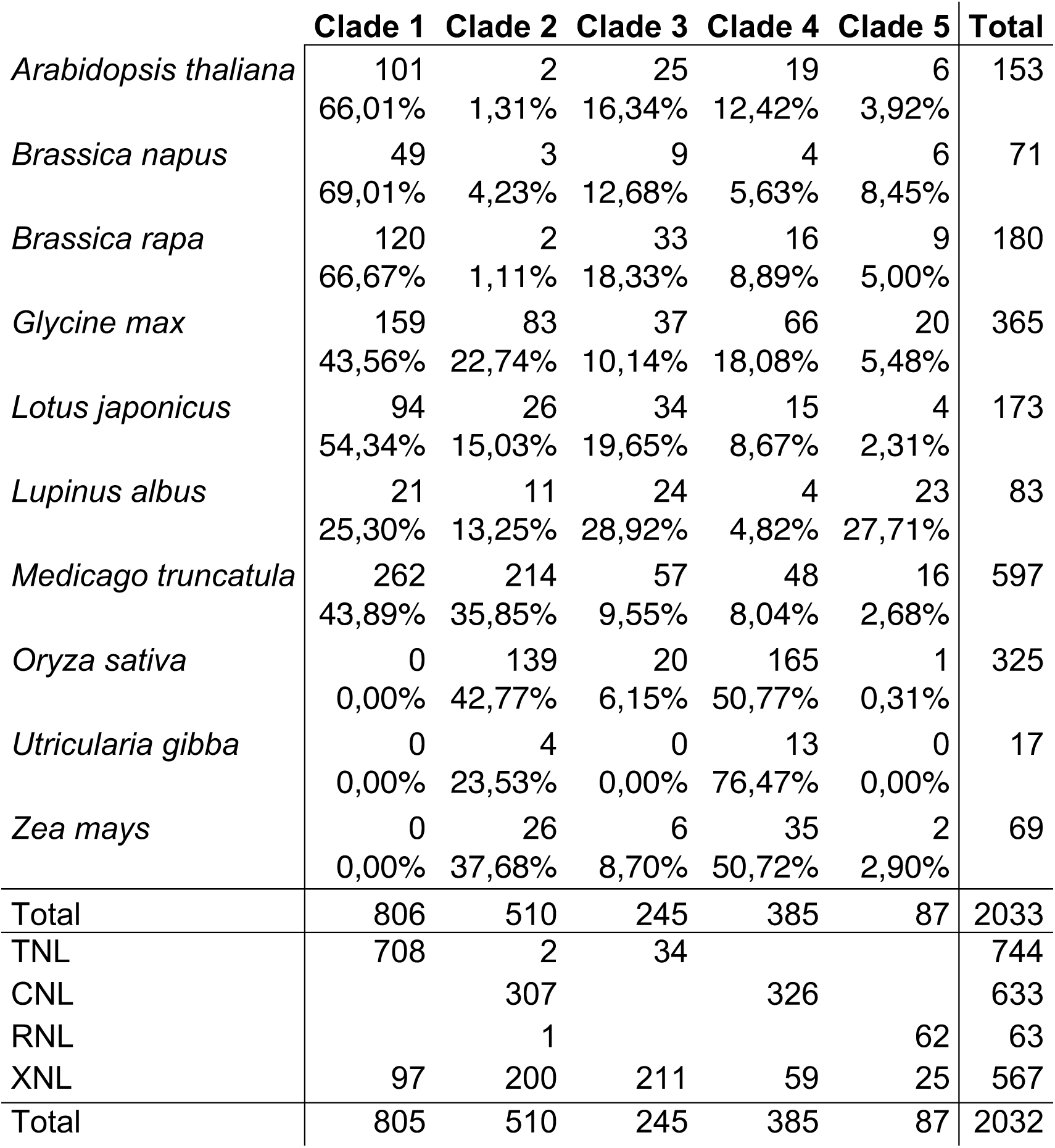
Clade distribution of NLR genes for the phylogenetic tree used in Figure 2B, including number of identified NLRs identified with at least one TIR (TNL), CC (CNL) or CC_R_ (RNL) domain, or none of the former three (XNL).

**Supplemental table 8.**
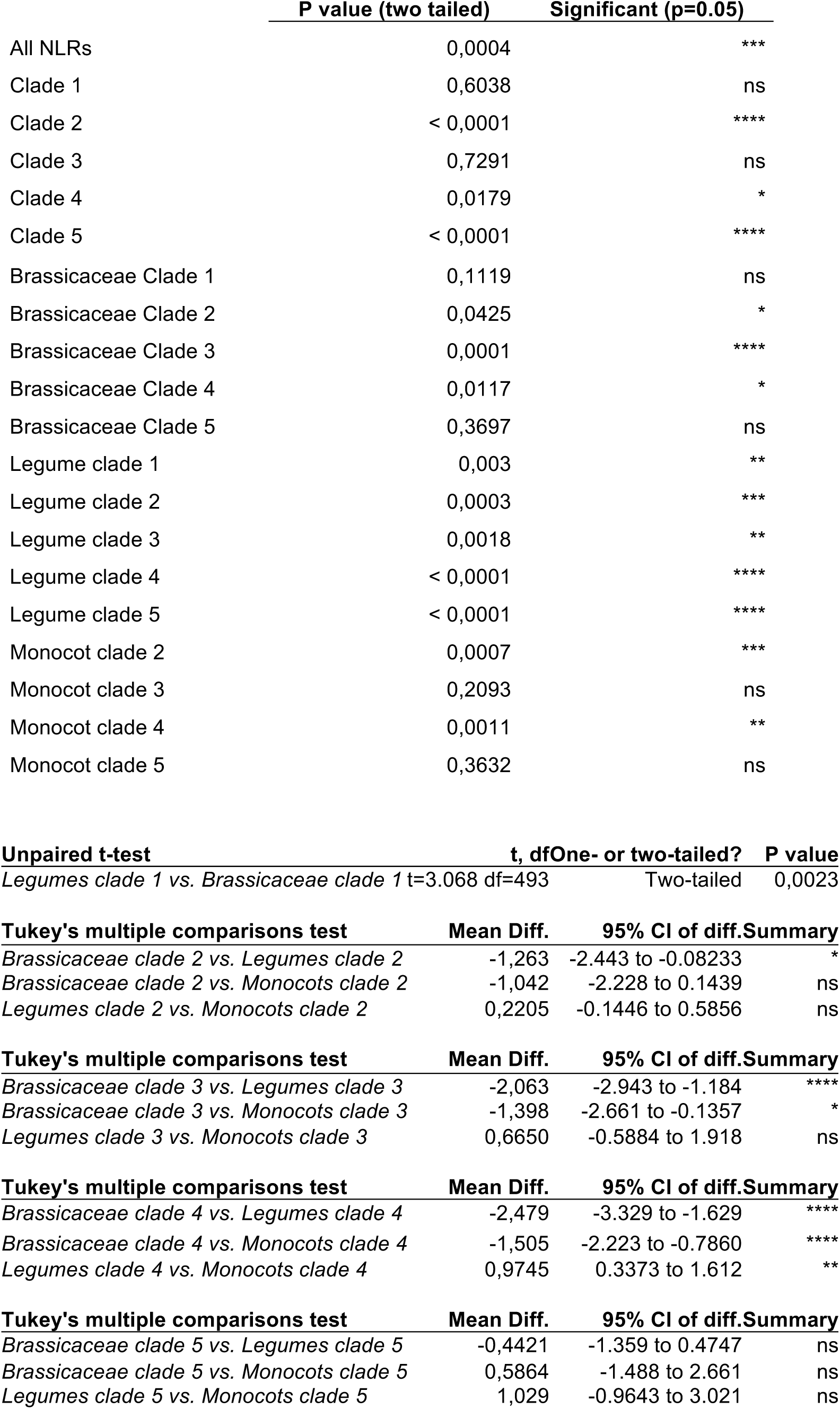
Cross-species clade comparison of normalized log_2_ root/shoot NLR gene expression ratios. ANOVA and Tukey’s multiple comparison test was used for calculation of *P*-values.

**Supplemental Table 9.** Clade distribution of NLR genes for the phylogenetic tree used in Figure 3B, including number of identified NLRs identified with at least one TIR (TNL), CC (CNL) or CC_R_ (RNL) domain, or none of the former three (XNL).

|  | Clade 1 | Clade 2 | Clade 3 | Clade 4 | Clade 5 | Total |
| --- | --- | --- | --- | --- | --- | --- |
| *Arabidopsis lyrata* | 37<br>54,41% | 3<br>4,41% | 20<br>29,41% | 6<br>8,82% | 2<br>2,94% | 68 |
| *Arabidopsis thaliana* | 101<br>66,01% | 3<br>1,96% | 24<br>15,69% | 19<br>12,42% | 6<br>3,92% | 153 |
| *Beta vulgaris* | 1<br>2,04% | 22<br>44,90% | 19<br>38,78% | 7<br>14,29% | 0<br>0,00% | 49 |
| *Brassica napus* | 49<br>69,01% | 6<br>8,45% | 9<br>12,68% | 1<br>1,41% | 6<br>8,45% | 71 |
| *Brassica rapa* | 121<br>66,85% | 3<br>1,66% | 32<br>17,68% | 16<br>8,84% | 9<br>4,97% | 181 |
| *Capsella rubella* | 16<br>34,04% | 2<br>4,26% | 20<br>42,55% | 6<br>12,77% | 3<br>6,38% | 47 |
| *Dianthus caryophyllus* | 1<br>1,85% | 31<br>57,41% | 10<br>18,52% | 12<br>22,22% | 0<br>0,00% | 54 |
| *Eutrema salsugineum* | 9<br>37,50% | 3<br>12,50% | 7<br>29,17% | 5<br>20,83% | 0<br>0,00% | 24 |
| *Glycine max* | 157<br>43,13% | 83<br>22,80% | 36<br>9,89% | 67<br>18,41% | 21<br>5,77% | 364 |
| *Lotus japonicus* | 93<br>54,39% | 24<br>14,04% | 34<br>19,88% | 16<br>9,36% | 4<br>2,34% | 171 |
| *Lupinus albus* | 21<br>25,00% | 11<br>13,10% | 24<br>28,57% | 5<br>5,95% | 23<br>27,38% | 84 |
| *Medicago truncatula* | 263<br>43,98% | 214<br>35,79% | 57<br>9,53% | 48<br>8,03% | 16<br>2,68% | 598 |
| *Nelumbo nucifera* | 2<br>3,13% | 34<br>53,13% | 8<br>12,50% | 20<br>31,25% | 0<br>0,00% | 64 |
| *Oryza sativa* | 0<br>0,00% | 137<br>42,55% | 17<br>5,28% | 167<br>51,86% | 1<br>0,31% | 322 |
| *Spirodela polyrhiza* | 0<br>0,00% | 22<br>34,92% | 23<br>36,51% | 18<br>28,57% | 0<br>0,00% | 63 |
| *Tarenaya hassleriana* | 20<br>40,82% | 3<br>6,12% | 23<br>46,94% | 3<br>6,12% | 0<br>0,00% | 49 |
| *Utricularia gibba* | 0<br>0,00% | 4<br>23,53% | 0<br>0,00% | 13<br>76,47% | 0<br>0,00% | 17 |
| *Zea mays* | 0<br>0,00% | 26<br>37,68% | 3<br>4,35% | 38<br>55,07% | 2<br>2,90% | 69 |
| **Total** | **891** | **631** | **366** | **467** | **93** | **2448** |
| TNL | 782 | 1 | 26 | 11 |  | 744 |
| CNL |  | 392 |  | 388 |  | 633 |
| RNL |  | 1 |  |  | 68 | 63 |
| XNL | 109 | 237 | 340 | 68 | 25 | 567 |
| **Total** | **891** | **631** | **366** | **467** | **93** | **2448** |

## References

Aarts N, M Metz, E Holub, Brian J Staskawicz, M J Daniels, and Jane E Parker. 1998. “Different Requirements for EDS1 and NDR1 by Disease Resistance Genes Define at Least Two R Gene-Mediated Signaling Pathways in *Arabidopsis*.” Proceedings of the National Academy of Sciences of the United States of America 95 (17): 10306–11.

Banba, Mari, Caroline Gutjahr, Akio Miyao, Hirohiko Hirochika, Uta Paszkowski, Hiroshi Kouchi, and Haruko Imaizumi-Anraku. 2008. “Divergence of Evolutionary Ways Among Common Sym Genes: CASTOR and CCaMK Show Functional Conservation Between Two Symbiosis Systems and Constitute the Root of a Common Signaling Pathway.” Plant and Cell Physiology 49 (11): 1659–71. doi:10.1093/pcp/pcn153.

Barker, David G, Sylvie Bianchi, François Blondon, Yvette Dattée, Gérard Duc, Sadi Essad, Pascal Flament, Philippe Gallusci, Gérard Génier, and Pierre Guy. 1990. “*Medicago Truncatula*, a Model Plant for Studying the Molecular Genetics of the Rhizobium-Legume Symbiosis.” Plant Molecular Biology Reporter 8 (8). Springer: 40–49. doi:10.1007/BF02668879.

Bartsev, Alexander V, William J Deakin, Nawal M Boukli, Crystal B McAlvin, Gary Stacey, Pia Malnoë, William J Broughton, and Christian Staehelin. 2004. “NopL, an Effector Protein of Rhizobium Sp. NGR234, Thwarts Activation of Plant Defense Reactions.” Plant Physiology 134 (2): 871–79. doi:10.1104/pp.103.031740.

Bellato C, H B Krishnan, T Cubo, F Temprano, and S G Pueppke. 1997. “The Soybean Cultivar Specificity Gene nolX Is Present, Expressed in a nodD-Dependent Manner, and of Symbiotic Significance in Cultivar-Nonspecific Strains of Rhizobium (Sinorhizobium) Fredii.” Microbiology (Reading, England) 143 (Pt 4) (April): 1381–88. doi:10.1099/00221287-143-4-1381.

Bonardi, Vera, Karen Cherkis, Marc T Nishimura, and Jeffery L Dangl. 2012. “A New Eye on NLR Proteins: Focused on Clarity or Diffused by Complexity?.” Current Opinion in Immunology 24 (1): 41–50. doi:10.1016/j.coi.2011.12.006.

Bonardi, Vera, Saijun Tang, Anna Stallmann, Melinda Roberts, Karen Cherkis, and Jeffery L Dangl. 2011. “Expanded Functions for a Family of Plant Intracellular Immune Receptors Beyond Specific Recognition of Pathogen Effectors.” Proceedings of the National Academy of Sciences of the United States of America 108 (39): 16463–68. doi:10.1073/pnas.1113726108.

Bulgarelli, Davide, Klaus Schlaeppi, Stijn Spaepen, Emiel ver Loren van Themaat, and Paul Schulze-Lefert. 2013. “Structure and Functions of the Bacterial Microbiota of Plants.” Annual Review of Plant Biology 64: 807–38. doi:10.1146/annurev-arplant-050312-120106.

Caplan, Jeffery L, Tadeusz Wroblewski, Richard W Michelmore, and Blake C Meyers. 2013. “The Role of TIR-NBS and TIR-X Proteins in Plant Basal Defense Responses.” Plant Physiology 162 (3): 1459–72. doi:10.1104/pp.113.219162.

Chen, Wei-Hua, and Martin J Lercher. 2009. “ColorTree: a Batch Customization Tool for Phylogenic Trees.” BMC Research Notes 2 (July): 155. doi:10.1186/1756-0500-2-155.

Deguchi, Yuichi, Mari Banba, Yoshikazu Shimoda, Svetlana A Chechetka, Ryota Suzuri, Yasuhiro Okusako, Yasuhiro Ooki, et al. 2007. “Transcriptome Profiling of *Lotus* Japonicus Roots During Arbuscular Mycorrhiza Development and Comparison with That of Nodulation.” DNA Research : an International Journal for Rapid Publication of Reports on Genes and Genomes 14 (3): 117–33. doi:10.1093/dnares/dsm014.

Delaux, Pierre-Marc, Kranthi Varala, Patrick P Edger, Gloria M Coruzzi, J Chris Pires, and Jean-Michel Ané. 2014. “Comparative Phylogenomics Uncovers the Impact of Symbiotic Associations on Host Genome Evolution.” PLoS Genetics 10 (7): e1004487. doi:10.1371/journal.pgen.1004487.

Dobin, Alexander, Carrie A Davis, Felix Schlesinger, Jorg Drenkow, Chris Zaleski, Sonali Jha, Philippe Batut, Mark Chaisson, and Thomas R Gingeras. 2013. “STAR: Ultrafast Universal RNA-Seq Aligner.” Bioinformatics (Oxford, England) 29 (1): 15–21. doi:10.1093/bioinformatics/bts635.

Doyle, J J. 1998. “Phylogenetic Perspectives on Nodulation: Evolving Views of Plants and Symbiotic Bacteria.” Trends in Plant Science. doi:10.1094/MPMI-05-11-0114.

Eddy, Sean R. 2011. “Accelerated Profile HMM Searches.” PLoS Computational Biology 7 (10): pe1002195. doi:10.1371/journal.pcbi.1002195.

Erb, Matthias, Claudia Lenk, Jörg Degenhardt, and Ted C J Turlings. 2009. “The Underestimated Role of Roots in Defense Against Leaf Attackers.” Trends in Plant Science 14 (12): 653–59. doi:10.1016/j.tplants.2009.08.006.

Falk, A, Bart J Feys, L N Frost, Jonathan D G Jones, M J Daniels, and Jane E Parker. 1999. “EDS1, an Essential Component of R Gene-Mediated Disease Resistance in *Arabidopsis* Has Homology to Eukaryotic Lipases.” Proceedings of the National Academy of Sciences of the United States of America 96 (6): 3292–97.

Fatima, Urooj, and Muthappa Senthil-Kumar. 2015. “Plant and Pathogen Nutrient Acquisition Strategies.” Frontiers in Plant Science 6: 750. doi:10.3389/fpls.2015.00750.

Fluhr, R. 2001. “Sentinels of Disease. Plant Resistance Genes.” Plant Physiology 127 (4): 1367–74.

Fu, Limin, Beifang Niu, Zhengwei Zhu, Sitao Wu, and Weizhong Li. 2012. “CD-HIT: Accelerated for Clustering the Next-Generation Sequencing Data.” Bioinformatics (Oxford, England) 28 (23): 3150–52. doi:10.1093/bioinformatics/bts565.

Gualtieri, G, and T Bisseling. 2000. “The Evolution of Nodulation.” Plant Molecular Biology 42 (1): 181–94.

Guillotin, Bruno, Jean-Malo Couzigou, and Jean-Philippe Combier. 2016. “NIN Is Involved in the Regulation of Arbuscular Mycorrhizal Symbiosis.” Frontiers in Plant Science 7: 1704. doi:10.3389/fpls.2016.01704.

Haas, Brian J, Alexie Papanicolaou, Moran Yassour, Manfred Grabherr, Philip D Blood, Joshua Bowden, Matthew Brian Couger, et al. 2013. “De Novo Transcript Sequence Reconstruction From RNA-Seq Using the Trinity Platform for Reference Generation and Analysis.” Nature Protocols 8 (8): 1494–1512. doi:10.1038/nprot.2013.084.

Handberg, Kurt, and Jens Stougaard. 1992. “*Lotus* Japonicus, an Autogamous, Diploid Legume Species for Classical and Molecular Genetics.” The Plant Journal : for Cell and Molecular Biology 2 (2). Wiley Online Library: 487–96.

Hofius, Daniel, Dimitrios I Tsitsigiannis, Jonathan D G Jones, and John Mundy. 2007. “Inducible Cell Death in Plant Immunity.” Seminars in Cancer Biology 17 (2): 166–87. doi:10.1016/j.semcancer.2006.12.001.

Hofius, Daniel, Torsten Schultz-Larsen, Jan Joensen, Dimitrios I Tsitsigiannis, Nikolaj H T Petersen, Ole Mattsson, Lise Bolt Jørgensen, Jonathan D G Jones, John Mundy, and Morten Petersen. 2009. “Autophagic Components Contribute to Hypersensitive Cell Death in *Arabidopsis*.” Cell 137 (4): 773–83. doi:10.1016/j.cell.2009.02.036.

Huson, Daniel H, and Celine Scornavacca. 2012. “Dendroscope 3: an Interactive Tool for Rooted Phylogenetic Trees and Networks.” Systematic Biology 61 (6): 1061–67. doi:10.1093/sysbio/sys062.

Høgslund, Niels, Simona Radutoiu, Lene Krusell, Vera Voroshilova, Matthew A Hannah, Nicolas Goffard, Diego H Sanchez, et al. 2009. “Dissection of Symbiosis and Organ Development by Integrated Transcriptome Analysis of *Lotus* Japonicus Mutant and Wild-Type Plants.” PloS One 4 (8): e6556. doi:10.1371/journal.pone.0006556.

Ibarra-Laclette, Enrique, Eric Lyons, Gustavo Hernandez-Guzman, Claudia Anahí Pérez-Torres, Lorenzo Carretero-Paulet, Tien-Hao Chang, Tianying Lan, et al. 2013. “Architecture and Evolution of a Minute Plant Genome.” Nature 498 (7452): 94–98. doi:10.1038/nature12132.

Ingle, Robert A. 2011. “Defence Responses of *Arabidopsis* Thaliana to Infection by Pseudomonas Syringae Are Regulated by the Circadian Clock.” PloS One 6 (10): e26968. doi:10.1371/journal.pone.0026968.

Jacob, Florence, Saskia Vernaldi, and Takaki Maekawa. 2013. “Evolution and Conservation of Plant NLR Functions.” Frontiers in Immunology 4: 297. doi:10.3389/fimmu.2013.00297.

Jiménez-Guerrero, Irene, Francisco Pérez-Montaño, JoséAntonio Monreal, Gail M Preston, Helen Fones, Blanca Vioque, Francisco Javier Ollero, and Francisco Javier López-Baena. 2015. “The Sinorhizobium (Ensifer) Fredii HH103 Type 3 Secretion System Suppresses Early Defense Responses to Effectively Nodulate Soybean.” Molecular Plant-Microbe Interactions : MPMI 28 (7): 790–99. doi:10.1094/MPMI-01-15-0020-R.

Jones, D A, and JDG Jones. 1997. “The Role of Leucine-Rich Repeat Proteins in Plant Defences.” Advances in Botanical Research.

Jones, Jonathan D G, and Jeffery L Dangl. 2006. “The Plant Immune System.” Nature 444 (7117). Nature Publishing Group: 323–29. doi:10.1038/nature05286.

Kalo, Peter, Cynthia Gleason, Anne Edwards, John Marsh, Raka M Mitra, Sibylle Hirsch, Júlia Jakab, et al. 2005. “Nodulation Signaling in Legumes Requires NSP2, a Member of the GRAS Family of Transcriptional Regulators.” Science (New York, NY) 308 (5729): 1786–89. doi:10.1126/science.1110951.

Kambara, Kumiko, Silvia Ardissone, Hajime Kobayashi, Maged M Saad, Olivier Schumpp, William J Broughton, and William J Deakin. 2009. “Rhizobia Utilize Pathogen-Like Effector Proteins During Symbiosis.” Molecular Microbiology 71 (1): 92–106. doi:10.1111/j.1365-2958.2008.06507.x.

Kawaharada Y, S Kelly, M Wibroe Nielsen, C T Hjuler, K Gysel, A Muszyński, R W Carlson, et al. 2015. “Receptor-Mediated Exopolysaccharide Perception Controls Bacterial Infection.” Nature 523 (7560): 308–12. doi:10.1038/nature14611.

Khan, Madiha, Rajagopal Subramaniam, and Darrell Desveaux. 2015. “Of Guards, Decoys, Baits and Traps: Pathogen Perception in Plants by Type III Effector Sensors.” Current Opinion in Microbiology 29 (November): 49–55. doi:10.1016/j.mib.2015.10.006.

Kim, Won-Seok, and Hari B Krishnan. 2014. “A nopA Deletion Mutant of Sinorhizobium Fredii USDA257, a Soybean Symbiont, Is Impaired in Nodulation.” Current Microbiology 68 (2): 239–46. doi:10.1007/S00284-013-0469-4.

Kouchi, Hiroshi, Haruko Imaizumi-Anraku, Makoto Hayashi, Tsuneo Hakoyama, Tomomi Nakagawa, Yosuke Umehara, Norio Suganuma, and Masayoshi Kawaguchi. 2010. “How Many Peas in a Pod? Legume Genes Responsible for Mutualistic Symbioses Underground.” Plant and Cell Physiology 51 (9): 1381–97. doi:10.1093/pcp/pcq107.

Kroj, Thomas, Emilie Chanclud, Corinne Michel Romiti, Xavier Grand, and Jean-Benoit Morel. 2016. “Integration of Decoy Domains Derived From Protein Targets of Pathogen Effectors Into Plant Immune Receptors Is Widespread.” The New Phytologist, February. doi:10.1111/nph.13869.

Kunze, Gernot, Cyril Zipfel, Silke Robatzek, Karsten Niehaus, Thomas Boller, and Georg Felix. 2004. “The N Terminus of Bacterial Elongation Factor Tu Elicits Innate Immunity in *Arabidopsis* Plants.” The Plant Cell 16 (12): 3496–3507. doi:10.1105/tpc.104.026765.

Le Fevre, Ruth, Edouard Evangelisti, Thomas Rey, and Sebastian Schornack. 2015. “Modulation of Host Cell Biology by Plant Pathogenic Microbes.” Annual Review of Cell and Developmental Biology, September. doi:10.1146/annurev-cellbio-102314-112502.

Lévy, Julien, Cécile Bres, Rene Geurts, Boulos Chalhoub, Olga Kulikova, Gérard Duc, Etienne-Pascal Journet, et al. 2004. “A Putative Ca2+ and Calmodulin-Dependent Protein Kinase Required for Bacterial and Fungal Symbioses.” Science (New York, NY) 303 (5662): 1361–64. doi:10.1126/science.1093038.

Limpens, Erik, Carolien Franken, Patrick Smit, Joost Willemse, Ton Bisseling, and Rene Geurts. 2003. “LysM Domain Receptor Kinases Regulating Rhizobial Nod Factor-Induced Infection.” Science (New York, NY) 302 (5645): 630–33. doi:10.1126/science.1090074.

Liu, Tzu-Yin, Teng-Kuei Huang, Shu-Yi Yang, Yu-Ting Hong, Sheng-Min Huang, Fu-Nien Wang, Su-Fen Chiang, Shang-Yueh Tsai, Wen-Chien Lu, and Tzyy-Jen Chiou. 2016. “Identification of Plant Vacuolar Transporters Mediating Phosphate Storage.” Nature Communications 7 (March): 11095. doi:10.1038/ncomms11095.

Long, S R. 1989. “Rhizobium-Legume Nodulation: Life Together in the Underground.” Cell 56 (2): 203–14.

Lukasik, Ewa, and Frank L W Takken. 2009. “STANDing Strong, Resistance Proteins Instigators of Plant Defence.” Current Opinion in Plant Biology 12 (4): 427–36. doi:10.1016/j.pbi.2009.03.001.

Madsen, Esben Bjørn, Lene Heegaard Madsen, Simona Radutoiu, Magdalena Olbryt, Magdalena Rakwalska, Krzysztof Szczyglowski, Shusei Sato, et al. 2003. “A Receptor Kinase Gene of the LysM Type Is Involved in Legume Perception of Rhizobial Signals.” Nature 425 (6958): 637–40. doi:10.1038/nature02045.

Madsen, Lene H, Leïla Tirichine, Anna Jurkiewicz, John T Sullivan, Anne B Heckmann, Anita S Bek, Clive W Ronson, Euan K James, and Jens Stougaard. 2010. “The Molecular Network Governing Nodule Organogenesis and Infection in the Model Legume *Lotus* Japonicus.” Nature Communications 1: 10. doi:10.1038/ncomms1009.

Marchler-Bauer, Aron, Shennan Lu, John B Anderson, Farideh Chitsaz, Myra K Derbyshire, Carol DeWeese-Scott, Jessica H Fong, et al. 2011. “CDD: a Conserved Domain Database for the Functional Annotation of Proteins.” Nucleic Acids Research 39 (Database issue): D225–29. doi:10.1093/nar/gkq1189.

Marquenet, Emélie, and Evelyne Richet. 2007. “How Integration of Positive and Negative Regulatory Signals by a STAND Signaling Protein Depends on ATP Hydrolysis.” Molecular Cell 28 (2): 187–99. doi:10.1016/j.molcel.2007.08.014.

Meinhardt L W, H B Krishnan, P A Balatti, and S G Pueppke. 1993. “Molecular Cloning and Characterization of a Sym Plasmid Locus That Regulates Cultivar-Specific Nodulation of Soybean by Rhizobium Fredii USDA257.” Molecular Microbiology 9 (1): 17–29.

Meyers B C, A W Dickerman, R W Michelmore, S Sivaramakrishnan, B W Sobral, and N D Young. 1999. “Plant Disease Resistance Genes Encode Members of an Ancient and Diverse Protein Family Within the Nucleotide-Binding Superfamily.” The Plant Journal : for Cell and Molecular Biology 20 (3): 317–32.

Meyers, Blake C, Alexander Kozik, Alyssa Griego, Hanhui Kuang, and Richard W Michelmore. 2003. “Genome-Wide Analysis of NBS-LRR-Encoding Genes in *Arabidopsis*.” The Plant Cell 15 (4): 809–34.

Meyers, Blake C, Michele Morgante, and Richard W Michelmore. 2002. “TIR-X and TIR-NBS Proteins: Two New Families Related to Disease Resistance TIR-NBS-LRR Proteins Encoded in *Arabidopsis* and Other Plant Genomes.” The Plant Journal : for Cell and Molecular Biology 32 (1): 77–92.

Millet, Yves A, Cristian H Danna, Nicole K Clay, Wisuwat Songnuan, Matthew D Simon, Danièle Werck-Reichhart, and Frederick M Ausubel. 2010. “Innate Immune Responses Activated in *Arabidopsis* Roots by Microbe-Associated Molecular Patterns.” The Plant Cell 22 (3): 973–90. doi:10.1105/tpc.109.069658.

Minh, Bui Quang, Minh Anh Thi Nguyen, and Arndt von Haeseler. 2013. “Ultrafast Approximation for Phylogenetic Bootstrap.” Molecular Biology and Evolution 30 (5): 1188–95. doi:10.1093/molbev/mst024.

Mun, Terry, Asger Bachmann, Vikas Gupta, Jens Stougaard, and Stig U Andersen. 2016. “*Lotus* Base: an Integrated Information Portal for the Model Legume *Lotus* Japonicus.” Scientific Reports 6 (December): 39447. doi:10.1038/srep39447.

Muthamilarasan, Mehanathan, and Manoj Prasad. 2013. “Plant Innate Immunity: an Updated Insight Into Defense Mechanism.” Journal of Biosciences 38 (2): 433–49.

Nguyen, Lam-Tung, Heiko A Schmidt, Arndt von Haeseler, and Bui Quang Minh. 2015. “IQ-TREE: a Fast and Effective Stochastic Algorithm for Estimating Maximum-Likelihood Phylogenies.” Molecular Biology and Evolution 32 (1): 268–74. doi:10.1093/molbev/msu300.

Nishimura, Marc T, and Jeffery L Dangl. 2010. “*Arabidopsis* and the Plant Immune System.” The Plant Journal : for Cell and Molecular Biology 61 (6): 1053–66. doi:10.1111/j.1365-313X.2010.04131.x.

Oldroyd, Giles E D. 2013. “Speak, Friend, and Enter: Signalling Systems That Promote Beneficial Symbiotic Associations in Plants.” Nature Reviews Microbiology 11 (4): 252–63. doi:10.1038/nrmicro2990.

Oldroyd, Giles E D, and J Allan Downie. 2006. “Nuclear Calcium Changes at the Core of Symbiosis Signalling.” Current Opinion in Plant Biology 9 (4): 351–57. doi:10.1016/j.pbi.2006.05.003.

Pan, Q, J Wendel, and R Fluhr. 2000. “Divergent Evolution of Plant NBS-LRR Resistance Gene Homologues in Dicot and Cereal Genomes.” Journal of Molecular Evolution 50 (3): 203–13. doi:10.1007/s002399910023.

Parniske, M. 2000. “Intracellular Accommodation of Microbes by Plants: a Common Developmental Program for Symbiosis and Disease?.” Current Opinion in Plant Biology 3 (4): 320–28. doi:10.1016/S1369-5266(00)00088-1.

Parniske, Martin. 2008. “Arbuscular Mycorrhiza: the Mother of Plant Root Endosymbioses.” Nature Reviews Microbiology 6 (10): 763–75. doi:10.1038/nrmicro1987.

Pel, Michiel J C, and Corné M J Pieterse. 2012. “Microbial Recognition and Evasion of Host Immunity.” Journal of Experimental Botany, October. doi:10.1093/jxb/ers262.

Radutoiu, Simona, Lene H Madsen, Esben B Madsen, Anna Jurkiewicz, Eigo Fukai, Esben M H Quistgaard, Anita S Albrektsen, Euan K James, Søren Thirup, and Jens Stougaard. 2007. “LysM Domains Mediate Lipochitin-Oligosaccharide Recognition and Nfr Genes Extend the Symbiotic Host Range.” The EMBO Journal 26 (17): 3923–35. doi:10.1038/sj.emboj.7601826.

Radutoiu, Simona, Lene Heegaard Madsen, Esben Bjørn Madsen, Hubert H Felle, Yosuke Umehara, Mette Grønlund, Shusei Sato, et al. 2003. “Plant Recognition of Symbiotic Bacteria Requires Two LysM Receptor-Like Kinases.” Nature 425 (6958): 585–92. doi:10.1038/nature02039.

Ranf, Stefanie, Nicolas Gisch, Milena Schäffer, Tina Illig, Lore Westphal, Yuriy A Knirel, Patricia M Sánchez-Carballo, et al. 2015. “A Lectin S-Domain Receptor Kinase Mediates Lipopolysaccharide Sensing in Arabidopsis Thaliana.” Nature Immunology, March. doi:10.1038/ni.3124.

Saraste, M, P R Sibbald, and A Wittinghofer. 1990. “The P-Loop-a Common Motif in ATP-and GTP-Binding Proteins.” Trends in Biochemical Sciences 15 (11): 430–34.

Schauser L, A Roussis, J Stiller, and J Stougaard. 1999. “A Plant Regulator Controlling Development of Symbiotic Root Nodules.” Nature 402 (6758): 191–95. doi:10.1038/46058.

Schmid, Markus, Timothy S Davison, Stefan R Henz, Utz J Pape, Monika Demar, Martin Vingron, Bernhard Schölkopf, Detlef Weigel, and Jan U Lohmann. 2005. “A Gene Expression Map of *Arabidopsis* Thaliana Development.” Nature Genetics 37 (5): 501–6. doi:10.1038/ng1543.

Shao, Zhu-Qing, Jia-Yu Xue, Ping Wu, Yan-Mei Zhang, Yue Wu, Yue-Yu Hang, Bin Wang, and Jian-Qun Chen. 2016. “Large-Scale Analyses of Angiosperm Nucleotide-Binding Site-Leucine-Rich Repeat (NBS-LRR) Genes Reveal Three Anciently Diverged Classes with Distinct Evolutionary Patterns.” Plant Physiology, February. doi:10.1104/pp.15.01487.

Sievers, Fabian, Andreas Wilm, David Dineen, Toby J Gibson, Kevin Karplus, Weizhong Li, Rodrigo Lopez, et al. 2011. “Fast, Scalable Generation of High-Quality Protein Multiple Sequence Alignments Using Clustal Omega.” Molecular Systems Biology 7 (October): 539. doi:10.1038/msb.2011.75.

Singh, Sylvia, and Martin Parniske. 2012. “Activation of Calcium- and Calmodulin-Dependent Protein Kinase (CCaMK), the Central Regulator of Plant Root Endosymbiosis.” Current Opinion in Plant Biology 15 (4): 444–53. doi:10.1016/j.pbi.2012.04.002.

Skorpil, Peter, Maged M Saad, Nawal M Boukli, Hajime Kobayashi, Florencia Ares-Orpel, William J Broughton, and William J Deakin. 2005. “NopP, a Phosphorylated Effector of Rhizobium Sp. Strain NGR234, Is a Major Determinant of Nodulation of the Tropical Legumes Flemingia Congesta and Tephrosia Vogelii.” Molecular Microbiology 57 (5): 1304–17. doi:10.1111/j.1365-2958.2005.04768.x.

Smit, Patrick, John Raedts, Vladimir Portyanko, Frédéric Debellé, Clare Gough, Ton Bisseling, and Rene Geurts. 2005. “NSP1 of the GRAS Protein Family Is Essential for Rhizobial Nod Factor-Induced Transcription.” Science (New York, NY) 308 (5729): 1789–91. doi:10.1126/science.1111025.

Smith, Sally E, and David J Read. 2010. Mycorrhizal Symbiosis. Academic Press.

Takken, Frank L W, and Aska Goverse. 2012. “How to Build a Pathogen Detector: Structural Basis of NB-LRR Function.” Current Opinion in Plant Biology 15 (4): 375–84. doi:10.1016/j.pbi.2012.05.001.

Takken, Frank Lw, Mario Albrecht, and Wladimir Il Tameling. 2006. “Resistance Proteins: Molecular Switches of Plant Defence.” Current Opinion in Plant Biology 9 (4): 383–90. doi:10.1016/j.pbi.2006.05.009.

Tang, Fang, Shengming Yang, Jinge Liu, and Hongyan Zhu. 2016. “Rj4, a Gene Controlling Nodulation Specificity in Soybeans, Encodes a Thaumatin-Like Protein but Not the One Previously Reported.” Plant Physiology 170 (1): 26–32. doi:10.1104/pp.15.01661.

Tanigaki, Yusuke, Kenji Ito, Yoshiyuki Obuchi, Akiko Kosaka, Katsuyuki T Yamato, Masahiro Okanami, Mikko T Lehtonen, Jari P T Valkonen, and Motomu Akita. 2014. “Physcomitrella Patens Has Kinase-LRR R Gene Homologs and Interacting Proteins.” PloS One 9 (4): e95118. doi:10.1371/journal.pone.0095118.

Tarr, D Ellen K, and Helen M Alexander. 2009. “TIR-NBS-LRR Genes Are Rare in Monocots: Evidence From Diverse Monocot Orders.” BMC Research Notes 2: 197. doi:10.1186/1756-0500-2-197.

Thordal-Christensen, Hans. 2003. “Fresh Insights Into Processes of Nonhost Resistance.” Current Opinion in Plant Biology 6 (4): 351–57.

Tirichine, Leïla, Haruko Imaizumi-Anraku, Satoko Yoshida, Yasuhiro Murakami, Lene H Madsen, Hiroki Miwa, Tomomi Nakagawa, et al. 2006. “Deregulation of a Ca2+/Calmodulin-Dependent Kinase Leads to Spontaneous Nodule Development.” Nature 441 (7097): 1153–56. doi:10.1038/nature04862.

Tsukui, Takahiro, Shima Eda, Takakazu Kaneko, Shusei Sato, Shin Okazaki, Kaori Kakizaki- Chiba, Manabu Itakura, et al. 2013. “The Type III Secretion System of Bradyrhizobium Japonicum USDA122 Mediates Symbiotic Incompatibility with Rj2 Soybean Plants.” Applied and Environmental Microbiology 79 (3): 1048–51. doi:10.1128/AEM.03297-12.

Tsurumaru, H, T Yamakawa, and M Tanaka. 2008. “Tn 5 Mutants of Bradyrhizobium Japonicum Is-1 with Altered Compatibility with Rj 2-Soybean Cultivars.” Soil Science & Plant …. doi:10.1111/j.1747-0765.2007.00225.x.

Van Dam, N M, TOG Tytgat, and J A Kirkegaard. 2009. “Root and Shoot Glucosinolates: a Comparison of Their Diversity, Function and Interactions in Natural and Managed Ecosystems.” Phytochemistry Reviews. doi:10.1007/S11101-008-9101-9.

van der Biezen, E A, and J D Jones. 1998. “The NB-ARC Domain: a Novel Signalling Motif Shared by Plant Resistance Gene Products and Regulators of Cell Death in Animals.” Current Biology : CB 8 (7): R226–27.

van Veen, Hans, Divya Vashisht, Melis Akman, Thomas Girke, Angelika Mustroph, Emilie Reinen, Sjon Hartman, et al. 2016. “Transcriptomes of Eight *Arabidopsis* Thaliana Accessions Reveal Core Conserved, Genotype- and Organ-Specific Responses to Flooding Stress.” Plant Physiology 172 (2): 668–89. doi:10.1104/pp.16.00472.

Vandenkoornhuyse, Philippe, Achim Quaiser, Marie Duhamel, Amandine Le Van, and Alexis Dufresne. 2015. “The Importance of the Microbiome of the Plant Holobiont.” The New Phytologist 206 (4): 1196–1206. doi:10.1111/nph.13312.

Verdier, Jérôme, Ivone Torres-Jerez, Mingyi Wang, Andry Andriankaja, Stacy N Allen, Ji He, Yuhong Tang, Jeremy D Murray, and Michael K Udvardi. 2013. “Establishment of the *Lotus* Japonicus Gene Expression Atlas (LjGEA) and Its Use to Explore Legume Seed Maturation.” The Plant Journal : for Cell and Molecular Biology 74 (2): 351–62. doi:10.1111/tpj.12119.

Viprey V, A Del Greco, W Golinowski, W J Broughton, and X Perret. 1998. “Symbiotic Implications of Type III Protein Secretion Machinery in Rhizobium.” Molecular Microbiology 28 (6): 1381–89.

Wang, Wei, Jinyoung Yang Barnaby, Yasuomi Tada, Hairi Li, Mahmut Tör, Daniela Caldelari, Dae-un Lee, Xiang-Dong Fu, and Xinnian Dong. 2011. “Timing of Plant Immune Responses by a Central Circadian Regulator.” Nature 470 (7332): 110–14. doi:10.1038/nature09766.

Xiao S, S Ellwood, O Calis, E Patrick, T Li, M Coleman, and J G Turner. 2001. “Broad-Spectrum Mildew Resistance in *Arabidopsis* Thaliana Mediated by RPW8.” Science (New York, NY) 291 (5501): 118–20. doi:10.1126/science.291.5501.118.

Xin, Da-Wei, Sha Liao, Zhi-Ping Xie, Dagmar R Hann, Lea Steinle, Thomas Boller, and Christian Staehelin. 2012. “Functional Analysis of NopM, a Novel E3 Ubiquitin Ligase (NEL) Domain Effector of Rhizobium Sp. Strain NGR234.” PLoS Pathogens 8 (5): e1002707. doi:10.1371/journal.ppat.1002707.

Xin, Xiu-Fang, and Sheng Yang He. 2013. “Pseudomonas Syringae Pv. Tomato DC3000: a Model Pathogen for Probing Disease Susceptibility and Hormone Signaling in Plants.” Annual Review of Phytopathology 51: 473–98. doi:10.1146/annurev-phyto-082712-102321.

Xue, Jia-Yu, Yue Wang, Ping Wu, Qiang Wang, Le-Tian Yang, Xiao-Han Pan, Bin Wang, and Jian-Qun Chen. 2012. “A Primary Survey on Bryophyte Species Reveals Two Novel Classes of Nucleotide-Binding Site (NBS) Genes.” PloS One 7 (5): e36700. doi:10.1371/journal.pone.0036700.

Yue, Jia-Xing, Blake C Meyers, Jian-Qun Chen, Dacheng Tian, and Sihai Yang. 2012. “Tracing the Origin and Evolutionary History of Plant Nucleotide-Binding Site-Leucine-Rich Repeat (NBS-LRR) Genes.” The New Phytologist 193 (4): 1049–63. doi:10.1111/j.1469-8137.2011.04006.x.

Zhang, Ling, Xue-Jiao Chen, Huang-Bin Lu, Zhi-Ping Xie, and Christian Staehelin. 2011. “Functional Analysis of the Type 3 Effector Nodulation Outer Protein L (NopL) From Rhizobium Sp. NGR234: Symbiotic Effects, Phosphorylation, and Interference with Mitogen-Activated Protein Kinase Signaling.” Journal of Biological Chemistry 286 (37): 32178–87. doi:10.1074/jbc.M111.265942.

## Supplemental references

Dash S, Van Hemert J, Hong L, Wise RP, Dickerson JA. PLEXdb: gene expression resources for plants and plant pathogens. Nucleic Acids Res. 2012 Jan;40(Database issue):D1194–201.

He J, Benedito VA, Wang M, Murray JD, Zhao PX, Tang Y, et al. The Medicago truncatula gene expression atlas web server. BMC Bioinformatics. 2009;10:441.

Høgslund N, Radutoiu S, Krusell L, Voroshilova V, Hannah MA, Goffard N, et al. Dissection of symbiosis and organ development by integrated transcriptome analysis of lotus japonicus mutant and wild-type plants. PLOS ONE. 2009;4(8):e6556.

Liu T-Y, Huang T-K, Yang S-Y, Hong Y-T, Huang S-M, Wang F-N, et al. Identification of plant vacuolar transporters mediating phosphate storage. Nat Commun. 2016 Mar 31;7:11095.

O'Rourke JA, Yang SS, Miller SS, Bucciarelli B, Liu J, Rydeen A, et al. An RNA-Seq transcriptome analysis of orthophosphate-deficient white lupin reveals novel insights into phosphorus acclimation in plants. Plant Physiol. 2013 Feb;161(2):705–24.

Schmid M, Davison TS, Henz SR, Pape UJ, Demar M, Vingron M, et al. A gene expression map of Arabidopsis thaliana development. Nat Genet. 2005 May;37(5):501–6.

Tong C, Wang X, Yu J, Wu J, Li W, Huang J, et al. Comprehensive analysis of RNA-seq data reveals the complexity of the transcriptome in Brassica rapa. BMC genomics. 2013 Oct 7;14:689.

van Veen H, Vashisht D, Akman M, Girke T, Mustroph A, Reinen E, et al. Transcriptomes of Eight Arabidopsis thaliana Accessions Reveal Core Conserved, Genotype- and Organ-Specific Responses to Flooding Stress. Plant Physiol. 2016 Oct;172(2):668–89.

Verdier J, Torres-Jerez I, Wang M, Andriankaja A, Allen SN, He J, et al. Establishment of the Lotus japonicus Gene Expression Atlas (LjGEA) and its use to explore legume seed maturation. Plant J. 2013 Apr;74(2):351–62.

Yong H-Y, Zou Z, Kok E-P, Kwan B-H, Chow K, Nasu S, et al. Comparative transcriptome analysis of leaves and roots in response to sudden increase in salinity in Brassica napus by RNA-seq. Biomed Res Int. 2014;2014:467395.

